# Structures of kinetic intermediate states of HIV-1 reverse transcriptase DNA synthesis

**DOI:** 10.1101/2023.12.18.572243

**Authors:** Sandra Vergara, Xiahong Zhou, Ulises Santiago, James F Conway, Nicolas Sluis-Cremer, Guillermo Calero

**Author notes:** Correspondence to: Guillermo Calero or Nicolas Sluis-Cremer.

## Abstract

Reverse transcription of the retroviral single-stranded RNA into double-stranded DNA is an integral step during HIV-1 replication, and reverse transcriptase (RT) is a primary target for antiviral therapy. Despite a wealth of structural information on RT, we lack critical insight into the intermediate kinetic states of DNA synthesis. Using catalytically active substrates, and a novel blot/diffusion cryo-electron microscopy approach, we captured 11 structures that define the substrate binding, reactant, transition and product states of dATP addition by RT at 1.9 to 2.4 Å resolution in the active site. Initial dATP binding to RT-template/primer complex involves a single Mg^2+^ (site B), and promotes partial closure of the active site pocket by a large conformational change in the β3-β4 loop in the Fingers domain, and formation of a negatively charged pocket where a second “drifting” Mg^2+^ can bind (site A). During the transition state, the α-phosphate oxygen from a previously unobserved dATP conformer aligns with the site A Mg^2+^ and the primer 3′-OH for nucleophilic attack. In the product state, we captured two substrate conformations in the active site: 1) dATP that had yet to be incorporated into the nascent DNA, and 2) an incorporated dAMP with the pyrophosphate leaving group coordinated by metal B and stabilized through H-bonds in the active site of RT. This study provides insights into a fundamental chemical reaction that impacts polymerase fidelity, nucleoside inhibitor drug design, and mechanisms of drug resistance.

---

Reverse transcription of the retroviral single-stranded RNA genome into double stranded DNA is an essential step in HIV-1 replication. Reverse transcriptase (RT), a multifunctional enzyme that encodes for RNA- and DNA-dependent DNA polymerase and ribonuclease H activities, catalyzes the entire reverse transcription reaction, and drugs that target this enzyme form the cornerstone of antiretroviral therapy^1–3^. RT exhibits an ordered sequential mechanism of DNA synthesis: template/primer (T/P) nucleic acid substrate binding to RT precedes dNTP binding and incorporation^4^. The rate-limiting step of the nucleotidyl-transfer reaction is a conformational change that precisely aligns the dNTP substrate in the active site to facilitate phosphodiester bond formation^5,6^. dNTP addition is metal dependent and is driven by two Mg^2+^ ions, which align substrates in the active site^7^. The Site A metal, which has only been observed in a few crystal structures of an RT-T/P-dNTP ternary complex^8–10^, likely facilitates phosphodiester bond formation by lowering the pKa of the primer 3’-OH, allowing proton release to activate the nucleophile^11^. The Site B metal is thought to stabilize the incoming dNTP by coordinating with non-bridging oxygens of the triphosphate, and may participate in pyrophosphate (PPi) release following chemistry^12,13^.

To date, all published structures of an RT-T/P-dNTP ternary complex utilized either a dideoxy-terminated T/P substrate or non-incorporable dNTP mimics^8,9,14–16^. These structures represent catalytically inert complexes that do not support nucleotide addition. As such, we lack insight into the kinetic intermediate states that: (i) facilitate alignment of the dNTP substrate in the RT active site and define the rate-limiting step of the catalytic reaction; (ii) the contribution that Mg^2+^ plays in positioning of the dNTP substrate and in phosphodiester bond formation; and (iii) the conformational changes and transition states in the active site that drive catalysis. To address these knowledge gaps, we devised *in situ* “diffusion/blot experiments” for time-resolved cryo-electron microscopy (EM) where different combinations of enzyme and substrates are applied on opposite sides of a grid allowing mixing of both reaction components by diffusion through the matrix of holes in the support film, prior to blotting and the subsequent plunge-freeze procedure (see Methods). Increasing the temperature of the reaction during the diffusion/blotting step provided additional means to control the reaction. Here, we report eleven structures (Extended Data Table 1) that define five unique intermediate states of nucleotide incorporation by HIV-1 RT at resolutions from 1.9 to 2.4 Å in the active site. These novel structures elucidate, for the first time, conformational changes in the active site of RT that facilitate dNTP incorporation. This work sheds light into a fundamental chemical reaction that impacts polymerase fidelity, nucleoside inhibitor drug design, and mechanisms of drug resistance.

## Structure of HIV-1 RT in complex with a DNA aptamer T/P substrate

We first solved the structure of HIV-1 RT bound to a 38-mer hairpin DNA aptamer T/P substrate (RT-T/P) with 2.3 Å resolution around the active site (Fig. 1a and Extended Data Fig. 1a). The 38- mer hairpin DNA aptamer T/P has a high affinity for RT and allows for nucleotide incorporation^17^. The aptamer contains two 2′-O-methylated nucleotides that were used as markers to monitor positioning of the T/P. The density of the DNA aptamer was well-resolved (Fig. 1b), including the two methyl groups that were unequivocally observed in the density map (Fig. 1c). Our cryo-EM structure of the RT-T/P complex has an RMSD on Cαs of 0.64 Å with a similar RT-aptamer crystal structure (PDB:ID 5D3G)^18^ (Extended Data Fig. 2a); however, density around the β3-β4 loop residues and the Fingers domain is well-defined in the cryo-EM map but not in the X-ray electron density map (Extended Data Fig. 2b). In the dNTP-free structure, the β3-β4 loop residues are located ∼11 Å away from the active site creating an “open pocket” for dNTP binding (Extended Data Fig. 1a, and Fig. 2e, see below). The electrostatic surface potential of the β3-β4 loop flanking the dNTP binding pocket is positively charged (Extended Data Fig. 2c. left panel) and could potentially interact with the negatively charged phosphate tail. Moreover, all key structural elements of the polymerase active site, including density of all residues, were visualized in the cryo-EM maps (see below).

**Fig. 1:**
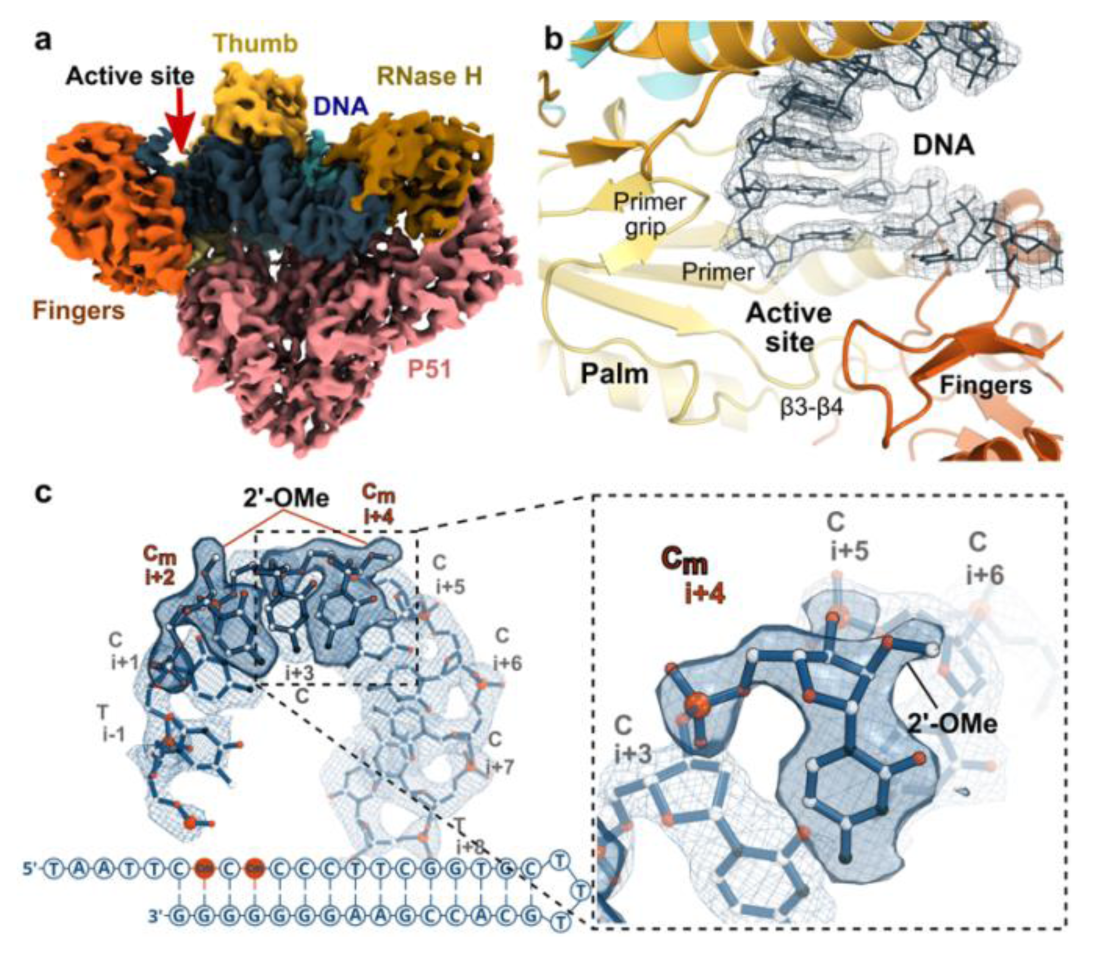
Cryo-EM structure of the dNTP-free RT-T/P complex. **a**, Cryo-EM density map for the RT-T/P complex colored by structural subdomains. **b**, Cartoon representation of the DNA polymerase active site of RT illustrating key protein motifs. The density for the T/P aptamer substrate is indicated in blue mesh with the corresponding model in sticks. **c**, Sequence of the 38-mer hairpin DNA aptamer T/P substrate used throughout this study. Nucleotides in positions +2 and +4 of the template strand labeled as Cm are 2’-O-methyl modified. Inset, detail of the density for the 2’-O-methyl modified nucleotide in position +4. Map sharpened at B-factor -80 Å^2^ and contoured at 6 σ.

**Fig. 2:**
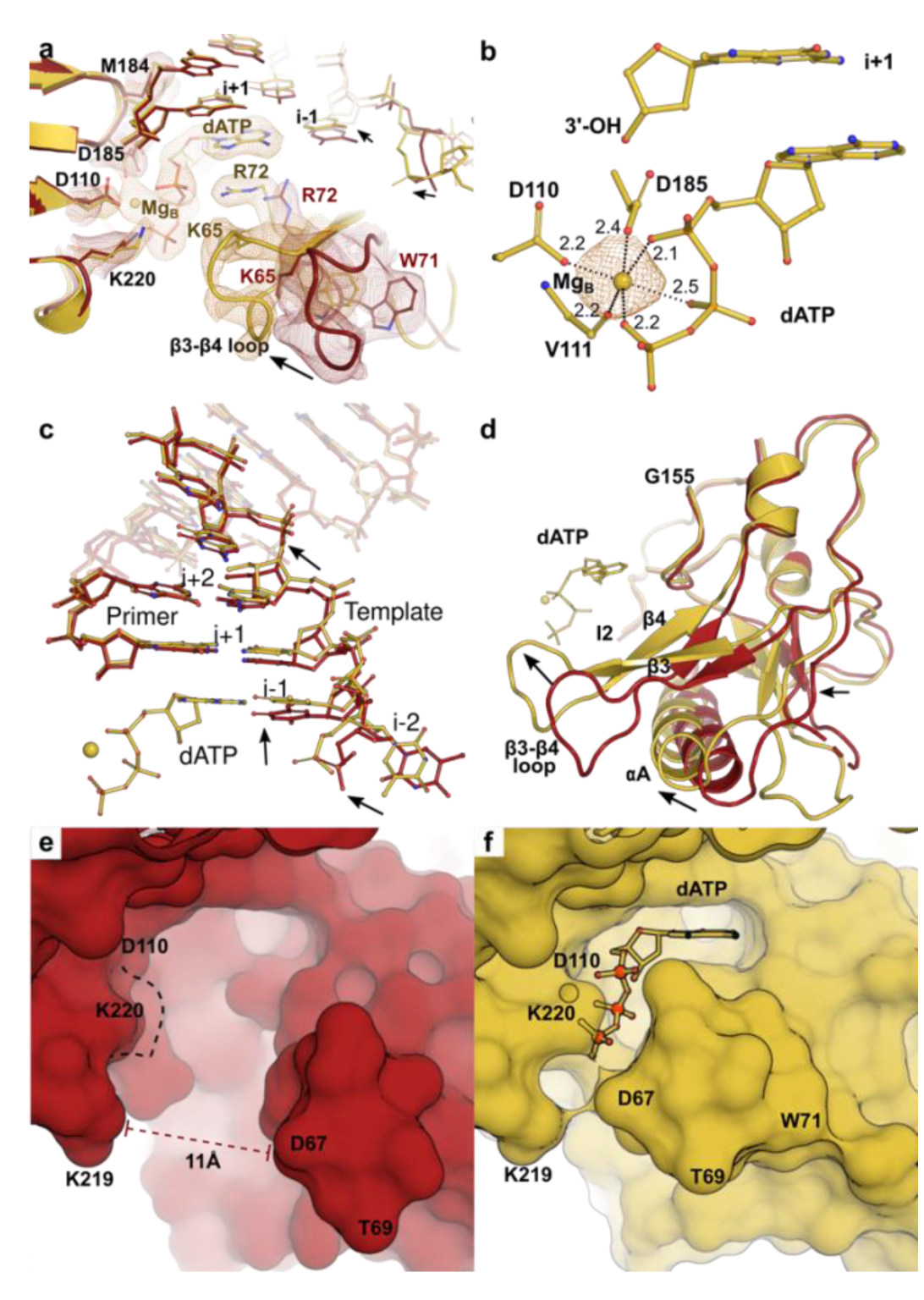
Cryo-EM structure of reactant state R-1 of the RT-T/P-dATP complex. **a**, Overlay of the structures of RT-T/P complex (red) and state R-1 of the RT-T/P-dATP complex (yellow) highlighting conformational changes in the RT active site that occur upon dATP binding (black arrows). Protein backbone displayed in cartoon and key residues displayed as ball-and-stick. Map for state R-1 is displayed in yellow mesh, with sharpening at B-factor -60 Å^2^ and contoured at 10 σ. Map for RT-T/P complex is displayed in red mesh, unsharpened, contoured at 7 σ. **b**, Site B Mg^2+^ coordination in state R-1 structure, yellow mesh contoured at 10 σ. Distances (in Å) are indicated next to dashed lines. **c,** Watson-Crick pairing of dATP with the templating (i-1) base, induces repositioning of the template strand (black arrows). **d**, Large-scale conformational changes of P66 Finger subdomain (residues 1–85 and 118–155) are observed (black arrows) upon dATP binding. **e**, Surface representation (shown in red) of the “Open” active site in the dNTP-free RT- T/P complex and **f**, the “Closed” active site of RT-T/P-dATP complex (shown in yellow) as a result of dATP binding (shown in ball-and-stick). Reposition of key residues D110 and K220 induces a tighter pocket in state R-1 evidenced by the extra bump region absent in RT-T/P (see panel **e**, dashed line).

## Structural basis of dNTP binding - Reactant state 1 of the RT-T/P-dATP complex

Using *in situ* diffusion/blot experiments, we successfully trapped an RT-T/P-dATP reactant state (R-1) by applying a solution of RT-T/P on one side of the grid and a solution of dATP-MgCl2 on the opposite side and immediately initiating the blotting/freezing procedure (see Methods). The overall structure of state R-1 of the RT-T/P-dATP ternary complex, at 2.1 Å resolution around the active site (Fig. 2a and Extended Data Fig. 1b), includes a dATP molecule in the binding pocket in the presence of a catalytically active 3′-OH primer; and a site B Mg^2+^ ion, coordinated by the α, β, and γ-phosphates, the backbone carbonyl of V111, and the carboxylic oxygens of D110 and D185 (Fig. 2b). Overlay of the RT-T/P and state R-1 structures shows an overall RMSD on Cαs of 0.52 Å. However, conformational changes can be observed in the aptamer templating base (Fig. 2c); the Fingers domain (RMSD 2.29 Å RT-T/P vs RT-T/P-dATP Fingers subdomain); the β3-β4 loop, which swings towards the substrate and closes the active site pocket (Fig. 2d); and active site residues D110 and K220 (Fig. 2e and f). In this way, the position of the β3-β4 loop away from the nucleotide binding site (in the RT-T/P structure) defines an “open” active site (Fig. 2e), while interactions of the β3-β4 loop with the incoming dATP define a “closed” active site (Fig. 2f). In the closed state, β3-β4 residue R72 stacks against the dATP base forming salt bridges with the closest α-phosphate oxygen while K65 forms a salt bridge with the γ-phosphate oxygen (Fig. 2a and Extended Data Fig. 3a). The positively charged β3-β4 loop (Extended Data Fig. 2c) together with conformational changes in active site residues (loop D110 to Y115 and K220) create a positive surface potential to bind the negatively charged phosphate tail (Extended Data Fig. 3b,c). Other residues, including D185, and those involved in dNTP binding stabilization and substrate selection, M184, Q151, Y115 and F116, also appear repositioned (Extended Data Fig. 3a).

## Reactant state 2 of the RT-T/P-dATP complex

A second reactant state (R-2) of the RT-T/P-dATP complex was trapped by applying a RT-T/P solution preincubated with dATP on one side of the grid and MgCl2 on the other side, and then immediately initiating the blotting/freezing procedure. The structure features 2.4 Å resolution around the active site (Extended Data Fig. 1c) with a RMSD on Cαs of 0.34 Å with respect to the state R-1 structure. The overall structure of state R-2 is almost identical to R-1, including site B Mg^2+^ ion. However, diffusion of Mg^2+^ to a pre-bound dATP enzyme allowed observation of strong, previously unreported density close to the side chain of D110, which we modeled as a Mg^2+^ ion (Fig. 3a-c). This putative Mg^2+^ does not correspond to the typical location of metal A reported for HIV-1 RT (Extended Data Fig. 4a)^8,9^; or the site C position observed in some DNA polymerases^19–21^, thus we designated it as site A* Mg^2+^. Resolving waters from ions in cryo-EM is an active area of analysis^22–24^. In this regard, the density at site A* was modeled as an ion rather than water based on the criteria described in the UnDowser procedure^22^ and the segmentation-guided water and ion modeling method^25^. These criteria consider that the distance of a water atom to a nearby polar atom is 2.8 ± ∼0.3 Å, whereas the distance of an ion to a charged/polar atom is 2.2 ± ∼0.3 Å. In the Coulomb potential map, site A* Mg^2+^ is located ∼2.3 Å from the carboxylic oxygen of D110.

**Fig. 3:**
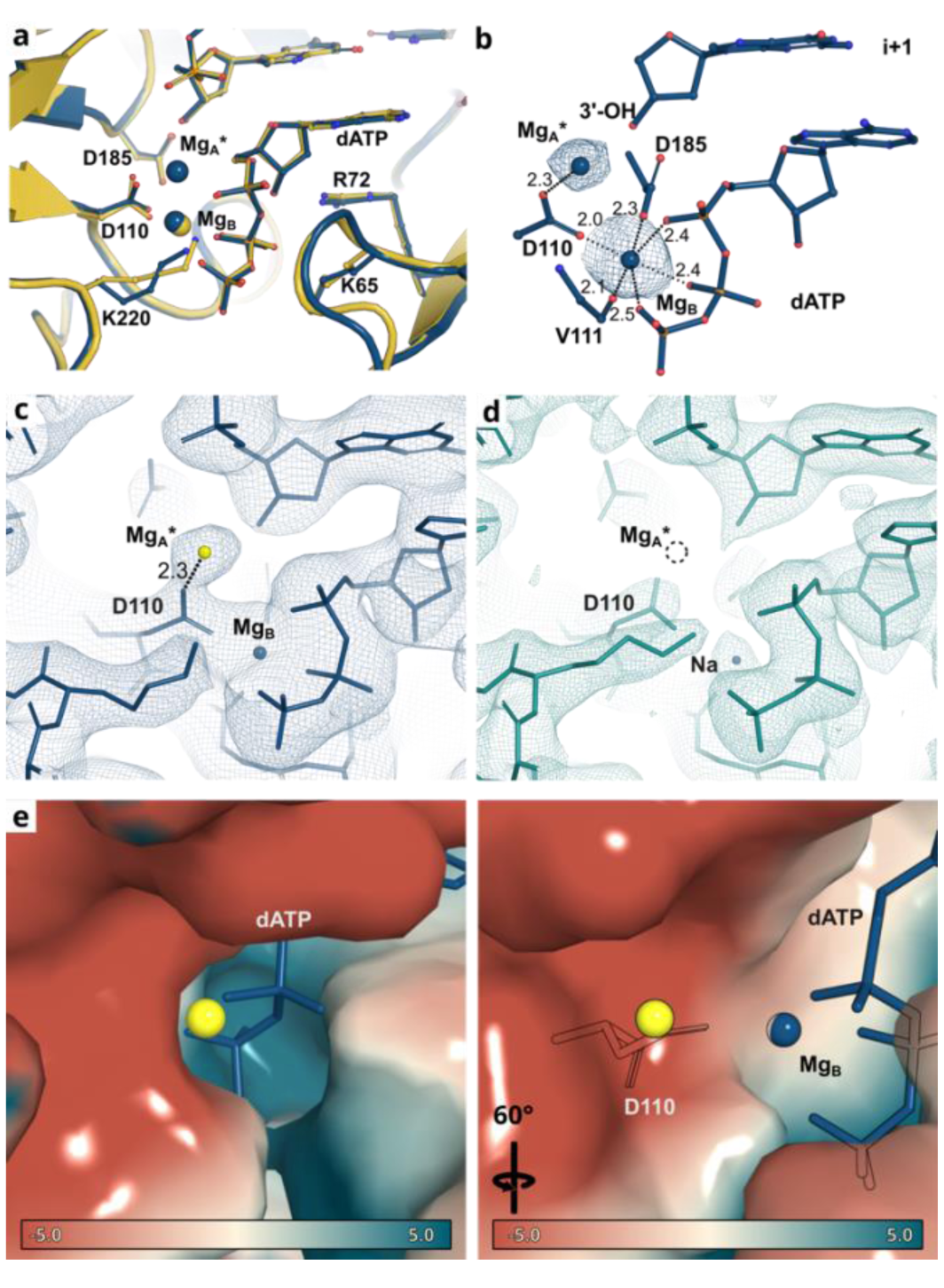
Cryo-EM structure of reactant state R-2 of the RT-T/P-dATP complex. **a**, Overlay of the active sites of states R-1 (yellow) and R-2 (blue) of the RT-T/P-dATP complex. The protein backbone is shown in cartoon representation; selected key residues are shown in ball- and-stick representation. **b**, Two Mg^2+^ sites, B and A*, are observed at state R-2. Map displayed as blue mesh (sharpening at B-factor = -80 Å^2^ and contoured at 11 σ). Coordination distances (in Å) are indicated next to dashed lines. **c**, Unsharpened density map for state R-2, contoured at 10 σ (blue mesh) showing density around site A* Mg^2+^ (yellow sphere); and **d**, unsharpened density map, contoured at 10 σ (teal mesh) for RT-T/P-dATP in the absence of MgCl2 showing no density at the A* position (dashed circle). **e**, Two views of the surface representation of RT-T/P-dATP in state R-2 colored according to electrostatic potential, from red (-5 kT/e, negatively charged) to blue (+5 kT/e, positively charged), showing Mg^2+^ A* positioned in a negatively charged pocket. Mg^2+^ ion site A* is colored as a yellow sphere.

A prior study reported that water molecules can display high variability in positions and contour shapes at different resolutions^23^. In contrast, ions appear in a similar position and shape in all maps, with ions still being observed at 3.1 Å resolution whereas waters are no longer detectable. The density at site A* was consistent with these criteria. First, the density consistently appeared at the same position in similar experimental conditions (Extended Data Fig. 4b) and the shape and position of the density were consistent in both half maps and the full map (Extended Data Fig. 4c); moreover, there was no remarkable change when the map was sharpened at different B-factors, contrary to the noise amplification effect of sharpening (Extended Data Fig. 4d). Second, a control experiment where RT was incubated with dATP in the absence of MgCl2 but with EDTA, to remove any trace Mg^2+^ contaminant (see Methods), showed no density (that could correspond to site A* Mg^2+^) near the side chain of D110 (Fig. 3d). A small density was observed at site B and it was modeled as a Na^+^ ion since its distance to the nearby phosphate oxygens ranged between 2.4-2.8 Å (which is within the experimental Na-O distance, 2.4 Å) ^26,27^. This experiment illustrates that a Mg^2+^ ion is not a prerequisite for dNTP binding, however, it is required for activity. Third, calculation of the surface electrostatic potential around site A* metal shows the presence of a negative potential pocket that would favor recruitment of positive charges (Fig. 3e).

## A transition state of the RT-T/P-dATP complex

With the aim of broadening our experimental conditions, we performed two additional simultaneous perturbations to state R-2 conditions: 1) we allowed sample diffusion for 8 seconds before blotting/freezing, and 2) we changed the Vitrobot chamber temperature from 22 ^°^C to 37 ^°^C during vitrification (Extended Data Fig. 1d). Under these conditions we captured a state (2.4 Å resolution around the active site) in which we observed the primer 3′-OH group moving closer tothe α-phosphate (Fig. 4a, black arrow); and the density for site A* Mg^2+^ shifting towards the 3’- OH and the α-phosphate (Fig. 4b). Intriguingly, two conformations of the dATP’s phosphate tail could be observed. The first one is similar to that of dATP in states R-1 and R-2, and we suggest that this is a ground state (dATP-1). However, we observed additional continuous density around the phosphate moiety that cannot be accounted for by only one dATP molecule (Fig. 4c and Extended Data Fig. 5a,b). Indeed, we modeled a second conformer of dATP (dATP-2). In this conformation the phosphate tail is rotated about the γ-dihedral bond, which determines the orientation of the phosphate tail in relation to the sugar moiety. The dATP-2 is within coordination distance of the site A* Mg^2+^ ion and repositions the α-phosphate within 4.1 Å from the primer 3′- OH (Fig. 4b). This conformation could represent a transition state (T8sec). Of note, these ATP conformers were observed in two individual experiments, highlighting that this alternative conformation is unlikely to be an artifact (Extended Data Fig. 5c).

**Fig. 4:**
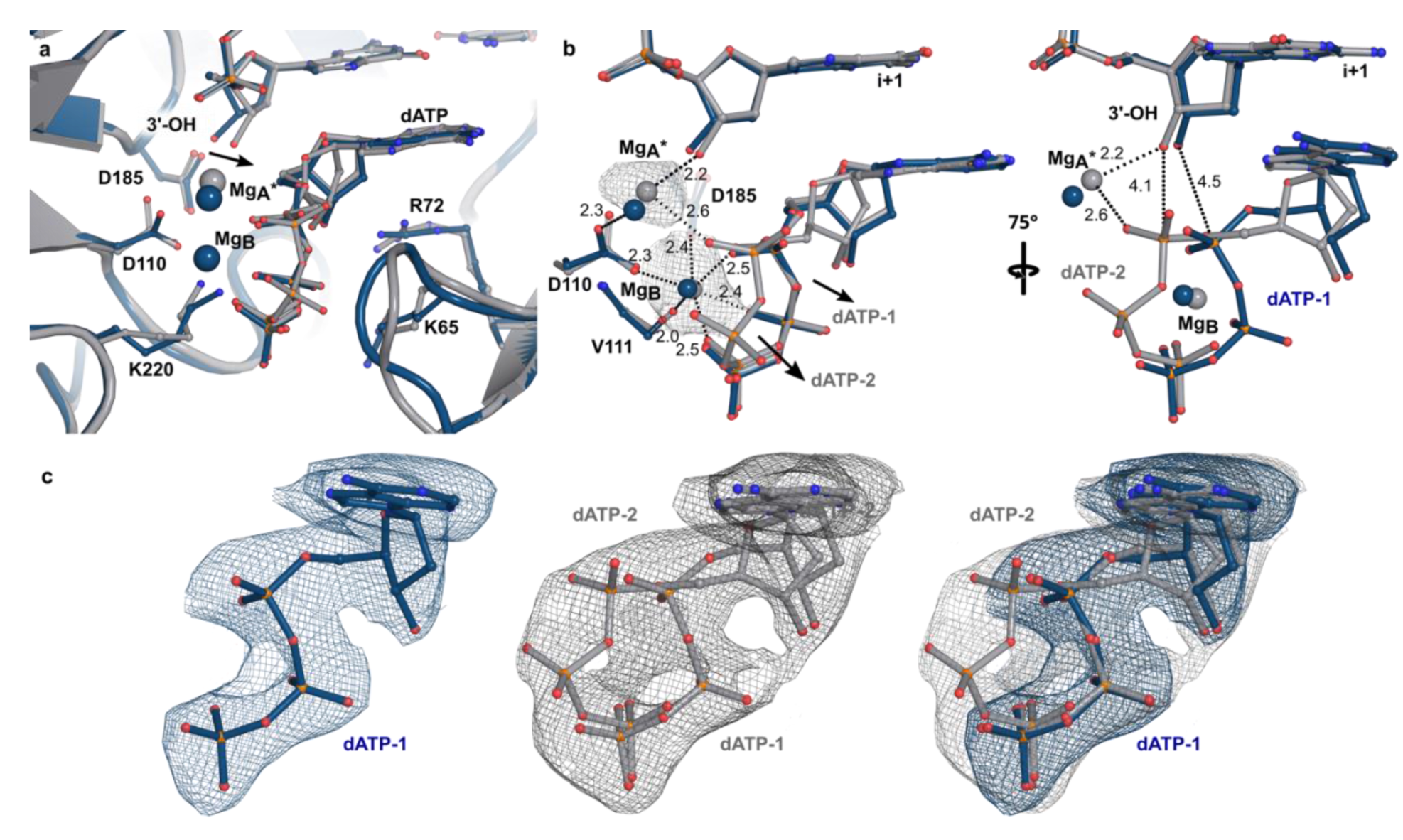
Cryo-EM structure of transition state of the RT-T/P-dATP complex. **a**, Overlay of the active sites of state R-2 (blue) and state T8sec (gray) of the RT-T/P-dATP complex. The protein backbone is shown in cartoon representation; selected key residues are shown in ball- and-stick representation. **b**, Coordination of Mg^2+^ at sites B and A* in the active site of RT by the two different dATP conformers (dATP-1 and dATP-2) that are observed in the transition state of the RT-T/P-dATP complex. The map around the Mg^2+^ ions at sites B and A* is shown as a grey mesh and is contoured at 8.5 σ (Map sharpened at B-factor -60 Å^2^). Coordination distances (in Å) are indicated next to dashed lines. **c**, Density around dATP-1 in state R-2 of the RT-T/P-dATP complex (blue mesh, left panel) and around dATP-1 and dATP-2 in state T8sec of the RT-T/P-dATP complex (gray mesh, middle panel). Overlay of the R-2 and T8sec state maps contoured at 13 σ and 8.5 σ, respectively, is shown in the right panel. Maps sharpened at B-factors -60 Å^2^ and -100 Å^2^ for states R-2 and T8sec, respectively.

## Product states of the RT-T/P-dATP complex

We trapped two product states of the RT-T/P-dATP complex by incubating a solution with all reactants (RT-T/P, dATP and MgCl2) for 2 and 6 minutes (P2min and P6min, respectively) at 37° C prior to grid preparation (active site resolution of 2.2 and 2.3 Å respectively; Extended Data Fig. 1e). Inspection of Coulomb potential maps for both states showed additional density for the 2′-O- methyl groups in the DNA aptamer at positions +3 and +5. For states R-1, R-2 and T8sec, we observed density at positions +2 and +4 as expected (Fig. 5a); however, the state P2min showed a mixture of partial occupancies at positions +2 and +3 and at +4 and +5 (Fig. 5b). Conversely, density in the state P6min had entirely shifted to the +3 and +5, indicating that one nucleotide was fully added and translocated (Fig. 5c). Further analysis of P2min and P6min maps highlighted further difference for these states. The P2min map showed that the position of the 3’-OH and site A* metal were similar to the state T8sec (with additional coordination from residue D110) thus showing a mixture of transition and product states (Extended Data Fig. 6a,b). The Coulomb potential map of P6min shows two conformations of the dATP in the binding pocket, which could be explained by a mixture of reactants and products with two chemical states of the substrate: one where density of the α-phosphate stretches toward the 3′-OH of the primer to form a phosphodiester bond and a scissile PPi (Fig. 5d); and a second one corresponding to an intact dATP, with density for the α-β-phosphate bond (Fig. 5e). To further corroborate the presence of the two dATP transition states in the active site pocket we overlayed the P6min map with that of the earliest state, R-1. Four major structural changes could be identified during addition (Fig. 5f): first, the primer 3′-OH swings 1.8 Å towards the incoming dATP; second, density for a phosphodiester bond emerges between the α-phosphate and the 3′-OH (black arrows); third, density weakens for the α- and β-phosphate bond (red arrows); and fourth, density at the phosphate tail is extended, indicative of PPi separation (green arrows). Indeed, increasing the density map threshold (to reveal only the highest scattering atoms) (Fig. 5g) shows that the α-phosphate center of density in state P6min shifts within bonding distance of the 3′-OH (1.9 Å vs 4.3 Å for state R-1), and that the distance between the α- and β- phosphate has increased beyond bonding length (3.7 Å vs a bonding distance of 3.0 Å in state R- 1). The latter suggests that a high proportion of dATP reactant molecules have been incorporated. The β-phosphate of the leaving PPi is coordinated by the site B metal while the γ-phosphate is stabilized through hydrogen bonding with K220, the amide nitrogen of D113 and β3-β4 loop K65 (Fig. 5h). In addition to Mg^2+^ ions at sites B and A*, density corresponding to the canonical site A metal^8,9^ could be observed; however, density was weak (around the noise level for the map) and could only be observed after map sharpening (Extended Data Fig. 7a). This was not surprising as Mg^2+^ at this canonical A site has been detected only in few crystal structures of an RT-T/P-dNTP ternary complex^8–10^; yet all of these structures had nucleic acid scaffolds featuring 3’ chain terminated primers. We next tested whether we could observe a Mg^2+^ ion at site A mimicking the conditions that allowed its observation in the original reports, i.e., employing a 3’ chain terminated primer (Extended Data Fig. 7b,c). Indeed, the cryo-EM density map shows the presence of density at the site A position, and no additional density at site A* (Extended Data Fig. 7d). Thus, the absence/presence of the 3’-OH primer could cause fluctuations in the location of the Mg^2+^ ion, suggesting that site A and site A* observed positions could relate to a single, oscillating Mg^2+^ ion. A control experiment with a non-hydrolyzable dNTP (dApCpp) showed a bound, yet frozen (not- added) substrate (Extended Data Fig. 6c). The structure exhibits well defined density for both sites B and A*. The latter metal is within coordination distance of D110, an α-phosphate oxygen and the primer’s 3′-OH, highlighting the high mobility of this Mg^+2^. Coordination of metal B showed small variations with the state R-2 structure, mainly due to differences in bond lengths (Extended Data Fig. 6d).

**Fig. 5:**
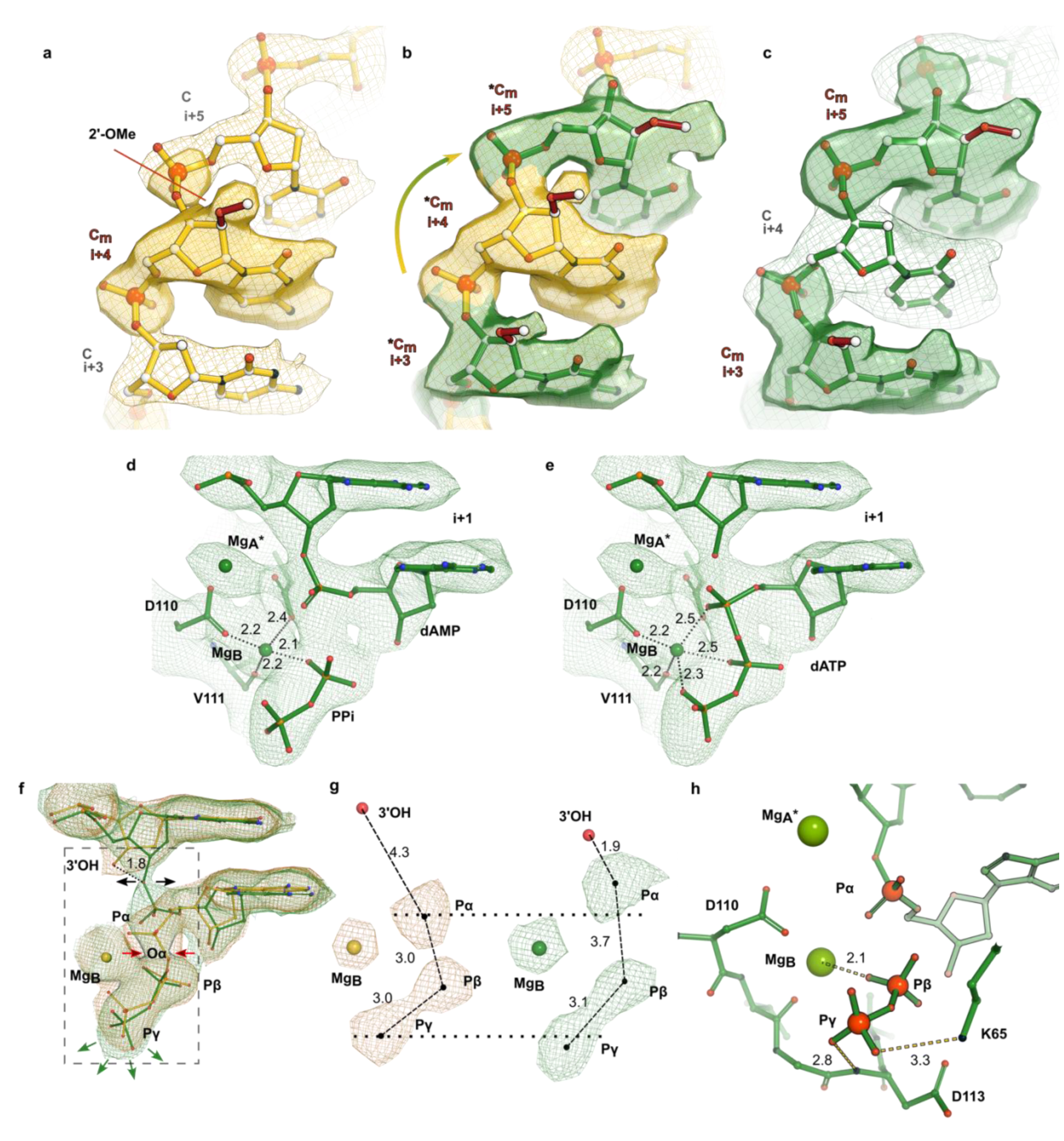
Cryo-EM structure of product states of the RT-T/P-dATP complex. Location of the 2′-O-methyl moiety in states (**a**) R-1, (**b**) P2min, and (**c**) P6min of the RT-T/P-dATP complex. **a**, In states R-1, R-2 and T8sec (R-2 and T8sec not displayed for clarity purpose) density (yellow mesh) for the 2’-OMe (Cm, red sticks) is observed in position +4 (and +2, not shown) and is clearly absent in positions +3 and +5 of the template strand (map displayed with sharpening of B-factor = -60 Å^2^ contoured at 12 σ). **b**, In state P2min density for 2’-OMe is observed in positions+2, +3, +4 and +5 as a mixture of translocated and non-translocated molecules. *Cm denotes partial occupancy of the 2’-OMe primer scaffold. Map density is colored with yellow mesh at position +4, and green mesh at positions +3 and +5 (contoured at 7.5 σ and sharpened at B-factor of -40 Å^2^). **c**, In P6min density for 2’-OMe (green mesh) is observed only in positions +3 and +5, indicating all molecules have translocated one position (map displayed with B-factor sharpening = -40 Å^2^ contoured at 8.5 σ). **d**,**e**, Ball-and-stick representation of the active site in state P6min with a mixture of (**d**) products (added dAMP and pyrophosphate) and (**e**) reactants (i.e., dATP). **f**, Overlay of density maps of states R-1 and P6min of the RT-T/P-dATP complex. Black arrows point to the growth of density for the newly formed phosphodiester bond; red arrows indicate thinning of density around O3A; green arrows point to stretched density for the leaving pyrophosphate (PPi) group (densities contoured at σ = 12.5 and σ = 9.5, for R-1 and P6min, respectively. Sharpened at B-factor of -60 Å^2^). **g**, Detail of density maps inside box in **f**, displayed at high threshold (σ = 35 and σ = 14, for state R-1 and state P6min, respectively. B-factor sharpening = -100 Å^2^). **h**, The PPi is stabilized by metal B and surrounding amino acids K65, D113 and K220 (not shown for clarity purpose). Distances are indicated next to dashed lines (in angstroms).

## Role for K220 in RT during nucleotide addition

K220 in RT is conserved across all HIV-1 isolates and has been implicated in a general acid catalysis mechanism of nucleotidyl transfer^28^. In all of our cryo-EM structures, K220 formed key interactions with the incoming dATP molecule that were not previously described in X-ray crystal structures. In state R-1, the incoming dATP promotes repositioning of the K220 side chain (Extended Data Fig. 8a) such that the Nζ moiety forms a salt bridge with the γ-phosphate of dATP. These interactions were maintained in state R-2 (not shown). In state T8sec, K220 interacts with the β-phosphate of the dATP-2 conformation (Extended Data Fig. 8b). We hypothesize that K220 together with the site A* Mg^2+^ induces and/or stabilizes the dATP-2 conformation. In state P6min, K220 displays two alternative conformations (Extended Data Fig. 8c). The first conformation is similar to state T8sec, while in the second conformation the Nζ points in the opposite direction which may be due to local conformational changes associated with the pyrophosphate group in the active site, as the orientation of the K220 side chain does not correspond to its orientation in the RT-T/P binary complex (Extended Data Fig. 8d). To probe the role of K220 in nucleotide incorporation, we introduced either leucine or methionine at residue 220 by side-directed mutagenesis (Extended Data Fig. 8e). Using transient kinetic analyses, we found that the K220L and K220M mutations in RT significantly diminished the rate of dATP incorporation (i.e., k_pol_) without substantially impacting the affinity of the nucleotide for the RT active site (i.e., K_d_), suggesting that K220 is involved in the catalytic event.

## Discussion

A two-metal-ion mechanism has been proposed for RT-mediated DNA synthesis as for other nucleic acid polymerases^7,13,29^. One metal ion forms a tight complex with the incoming dNTP (Mg^2+^ binding site B) via coordination with the non-bridging oxygens of all three phosphates, while the second (Mg^2+^ binding site A) binds to the closed state of the RT-T/P-(Mg^2+^) dNTP complex and reduces the pKa of the primer 3’-OH group, thereby activating the nucleophile and bringing it close to the α-phosphate of the incoming dNTP. The coordinated actions of the two metal ions, water molecules, and surrounding acidic residues are thought to stabilize the transition state. Here, using diffusion/blot experiments we captured RT-T/P-dATP-(Mg^2+^) structures encompassing substrate binding, and reactant, transition and product states. These structures reveal novel aspects of the chemistry of nucleotide addition.

During the transition from substrate binding to state R-1, the Fingers domain and the active site pocket undergo significant re-arrangements upon dNTP binding (Fig. 2e,f and Extended Data Fig. 3b,c). These changes include a rigid body rotation that propels the β3-β4 loop towards the dNTP resulting in partial closure of the active site pocket. This swing motion could be driven by a combination of two factors: 1) dNTP Watson-Crick base-pairing of the templating nucleotide pulling the template strand which lies right below it (Fig. 2a,c); and 2) electrostatic interactions between the positively charged β3-β4 loop and the dNTP (state R-1) (Extended Data Fig. 2c). Notably, our data show that in state R-1, site B Mg^2+^ (but not site A* metal) arrives first, and coordinates all three oxygen atoms of the phosphate tail; it is possible that this metal could be carried into the binding site pre-bound to the dNTP as the *K_d_* for Mg^2+^ binding to nucleotide is 29 μM^7^. Conversely, the Mg^2+^-free structure shows that a monovalent (sodium) cation can trigger dNTP binding and active site closure (Fig. 3d); yet, the enzyme is inactive.

Progression of state R-1 into R-2 involves binding of site A metal to a previously undescribed metal ion binding site in RT, site A* (Figs. 3 and 4). Structures for many classes of DNA polymerases have clearly defined density for Mg^2+^ sites A and B but, notably, a third Mg^2+^ ion has been observed for human DNA Pol η^19,30^, Pol β^20^ and Pol µ^21^. Binding of the third Mg^2+^ ion in the DNA Pol η, Pol β and Pol µ structures was transient and occurred during phosphodiester bond formation where it has been shown to stabilize the leaving pyrophosphate group^19,31^. Our cryo-EM structure shows fundamental differences between the active sites of DNA polymerases and HIV-1 RT. Structural overlay of DNA Pol η with RT shows a third Mg^2+^ ion across site A and B metals (Extended Data Fig. 7e). On the other hand, the tightly closed pocket (Fig. 2f and Extended Data Fig. 3c) observed in RT sterically hinders potential binding of a third Mg^2+^ (metal C) at a homologous position. However, a small negatively-charged pocket forms in the neighborhood of the aspartic acid triad, Asp110, Asp185, Asp186 and the 3′-OH of the primer template (Extended Data Fig. 9a). This pocket can accommodate binding of site A* metal at different positions. The structural data presented here suggest the possibility that metal A* is not fixed, hence varying experimental conditions such as presence or absence of the 3′-OH cause fluctuations in its position (Extended Data Fig. 9b); nonetheless, given its proximity to the a- phosphate and the 3′-OH, it would still play a fundamental role during catalysis.

Evolution of state R-2 into T8sec required a “boost” of activation energy; this transition state was achieved with samples exposed to temperatures between 30-37° C immediately before plunge-freezing. The structure for the state T8sec shows density in the Coulomb potential map that can accommodate two dATP conformers (Extended Data Fig. 5). The α-phosphate of dATP-2 moves within ∼4 Å of the 3′-OH primer and merges into the coordination sphere of the Mg^2+^ site A*. This observation is unique among all previous structural (and time resolved) work on DNA/RNA polymerases since high temperature data collection could not be achieved for X-ray crystallographic experiments.

The state P2min evolves into P6min (Fig. 5a-c). The former features a mixture of transient (i.e., position and coordination of metal A*) and product states with a population of non-translocated and translocated particles (i.e., partial occupancies of the 2′-O-methylated dCMPs at positions +2, +3 +4 and +5). Conversely, in P6min most or all particles have translocated one position. Moreover, state P6min captures two conformers of the dATP in the binding pocket: 1) a pre-translocated dATP, and 2) an inserted dAMP forming a phosphodiester bond with the primer and the released PPi coordinated by metal B and stabilized through H-bonds with the surrounding residues.

The structures presented here shed light on the chemistry of nucleotide addition, namely, the role of metal A* to drive the chemical reaction. A crucial event during nucleotide addition is the transient deprotonation of the 3′-OH for nucleophilic attack of the α-phosphate^32–34^; this event could be aspartate- or water-as-base-mediated^33^. In the former, RT’s D185 could act as the general base to deprotonate the 3′-OH; however, molecular simulations of aspartic acid residues (in DNA polymerase η) suggested that aspartate-deprotonation could be energetically unfavorable. In the water-as-base mechanism, which is energetically favored, a water molecule, originating from a Mg^2+^-coordinated water, could deprotonate de 3′-OH and thus trigger phosphoryl transfer. A water molecule coordinated by site A* metal may well be responsible for such nucleophilic attack. Our data shows the presence of two metals in HIV-1 RT’s active site, metals B and A*. Metal B, is unlikely to be involved since it is fully coordinated by three dNTP phosphate oxygens and three neighboring residues (Fig. 2b); conversely, given that site A* Mg^2+^ locates next to the α-phosphate and has low (residue) coordination number, it thus might provide the water molecule involved in 3′-OH deprotonation. Moreover, this event may be possibly triggered during site A* coordination by the dATP-2 conformer (Fig. 4b,c).

Intriguingly, no residue acting as a general acid (i.e., donating a proton) to the leaving pyrophosphate has been identified for HIV-1 RT; the structural and biochemical data presented here suggest that K220 may perhaps be this residue. Hence, it is possible that a water molecule coordinated by site A* metal could function as the general base, and K220 could function as the general acid^28^.

## A nucleotide addition cycle

The transition states suggest a structural pathway to nucleotide addition, i.e., a “structural dNTP addition cycle” (Fig. 6 and Supplementary Video 1). 1) In the RT-T/P binary complex, the β3-β4 loop is in the open conformation with an exposed templating nucleotide (i-1). 2) dNTP binds in the active site pocket. 3) Watson-Crick pairing of the incoming dNTP triggers sliding of the DNA/RNA hybrid template and closing of the binding pocket by the positively charged β3-β4 loop. 4) dNTP in the active site pocket attracts site A* metal. 5) Coordination of site A* metal elicits a pre-catalytic conformation. 6) Concurrence of the primer 3′-OH, the α-phosphate and site A* metal trigger phosphodiester bond formation and PPi release. 7) The DNA/RNA hybrid translocation swings the β3-β4 loop towards its open conformation.

**Fig. 6:**
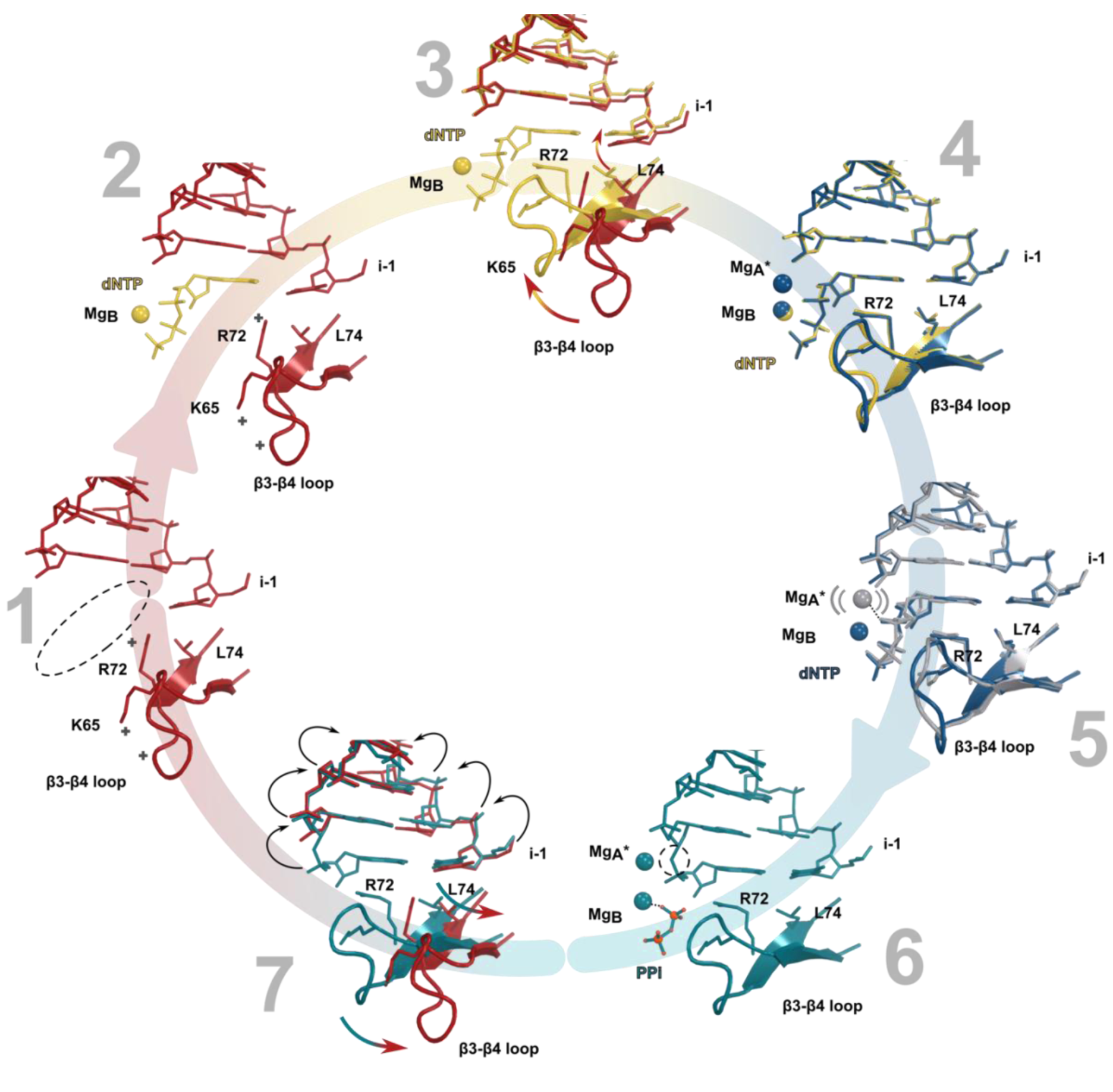
Proposed nucleotide addition cycle. Starting with the dNTP-free RT-T/P complex (red), we observed five unique intermediate states (R-1, R-2, T8sec, P2min, P6min) that suggests a possible cycle for nucleotide addition. See text for a detailed description.

The reactant, transition and product states presented here help to define the conformational changes that facilitate nucleotide incorporation. Collectively, this study provides insights into a fundamental chemical reaction that impacts polymerase fidelity, nucleoside inhibitor drug design, and mechanisms of drug resistance.

## Methods

### RT Expression and purification

The expression plasmid RT152A was obtained as a kind gift from Professor Eddy Arnold (Rutgers University). The protein was expressed in BL21(DE3) with 0.5 mM isopropyl β-d-1- thiogalactopyranoside at 16 °C overnight. Cells were lysed by sonication in 20 mM Tris pH 8.0, 150 mM NaCl, 5% Glycerol. After centrifugation at 18000 rpm for 40 min, the supernant was loaded to Nickel beads (HisTrap FF, Cytiva) and eluted with imidazole gradient from 0 mM to 500 mM. Collected fractions were analyzed with SDS-PAGE, and selected peak fractions were concentrated and loaded onto a heparin column (HiTrap heparin HP, Cytiva). Eluted fraction was concentrated and loaded to Superdex 200 10/300 GL column equilibrated with 20 mM Hepes pH 7.5, 100 mM NaCl, 5% Glycerol. The peak fractions were concentrated to 10 mg/mL, aliquoted and stored at -80°C.

### Preparation of the RT-T/P binary complex

The 38-mer DNA-hairpin T/P aptamer used in this study (5’- TAATTCCmCCmCCCTTCGGTGCTTTGCACCGAAGGGGGGG-3’) was purchased from Integrated DNA Technologies (Coralville, IA) (Fig. 1c). The RT-T/P was prepared by mixing RT with the T/P at a molar ratio of 1:1.3. The mixture was set on ice for 30 min before loading onto a Superdex 200 5/150 GL column equilibrated with 25 mM Tris-HCl pH 7.5, 100 mM NaCl, to remove the excess of DNA. The RT-T/P peak fraction was used for cryo-EM grid preparation.

### Cryo-EM grid preparation

We prepared cryo-EM grids with slightly different protocols for the eleven structures reported here (Extended Data Table 1). UltrAuFoil R1.2/1.3 300-mesh grids (Quantifoil Micro Tools GmbH, Großlöbichau, Germany) were freshly glow-discharged for 60 s and used immediately. If a solution was applied on both sides of the grid, each side of the grid was glow-discharged. Grids were vitrified with liquid ethane using a Vitrobot Mark IV (Thermo Fischer Scientific, USA) set at 22 °C (except for transition state T8sec), 100 % humidity, and a blotting time of 3 s. The final concentration of the liquid drop on the grid was 2.1 μM RT-T/P, 0.4 mM dATP, and 1 mM MgCl2. Solutions were kept on ice unless stated otherwise. For reactant states R-1 (data set 2) and R-2 (data sets 3 and 4) and transition states T8sec (data set 5) and T’8sec (data set 6) we performed “diffusion/blot experiments” where two separate solutions are applied on the opposite sides of the cryo-EM grid and mixing occurs on the grid by diffusion through the 1.2 µm diameter holes in the UltrAuFoil support (Fig. S1 and Extended Data Fig. 1). Application of the second solution starts the RT reaction. For state R-1, 2 μl of RT-T/P (4.2 μM) were applied on one side of the grid, and 2 μl of the second solution (0.8 mM dATP, 2 mM MgCl2) were applied to the opposite side of the grid, and the blotting/freezing cycle was immediately started (∼4 seconds of diffusion during blotting/freezing). For state R-2, 2 μl of RT-T/P preincubated with dATP (5 minutes, no MgCl2) were applied on the first side of the grid and 2 μl of solution containing MgCl2 were applied on the opposite side (blotting/freezing immediately). For state T8sec (data set 5) the Vitrobot and reactant solutions were pre-equilibrated at 37 °C (or 30 °C, for T’8sec data set 6) before solutions were applied to the grid as described for state R-2, but with a delay of 8 s before the blotting/freezing cycle was started. For the remaining cases (data sets 1, and 7 through 11), regular grid preparation protocols were followed, with 3 μl of sample applied to one side of the grid and the blotting/freezing cycle performed immediately. In the case of states P2min (data set 7) and P6min (data set 8), all components of the reaction were incubated in solution at 37 °C on a heat block for 2 minutes or 6 minutes, respectively, and then used immediately for grid preparation. For data set 9, the non-hydrolyzable dATP analogue dApCpp (2’-Deoxyadenosine-5’-[(α,β)- methyleno]triphosphate, Sodium salt, Jena Bioscience, Germany) was incubated with RT-T/P and MgCl2 at 37 °C for 15 minutes. To obtain the structure of RT-T/P-dATP in the absence of MgCl2 (data set 10), RT-T/P purification buffer was supplemented with 1 mM EDTA; dATP was preincubated for 1 h with EDTA (4mM dATP, 40mM EDTA) and subsequently all components (RT-T/P-dATP) were incubated 10 minutes at room temperature (final concentration 0.4mM dATP, 4mM EDTA). For the structure of the 3’ chain-terminated primer (data set 11), RT-T/P was prepared as described above with a ddCMP-terminated DNA aptamer (5’- TAATTGCmCCmCCCTTCGGTGCTTTGCACCGAAGGGGGG-ddC-3’) (Integrated DNA Technologies). In this case, the final concentration of MgCl2 was increased to 13 mM.

### Data collection

Cryo-EM data collection was performed using EPU software on a Titan Krios 3Gi transmission electron microscope operated at 300 kV (Thermo Fisher Scientific, USA). Movies were collected on a Falcon 3 direct electron detecting CMOS camera at a nominal magnification of 96,000 x corresponding to a calibrated pixel size of 0.825 Å (Data sets 1 to 6, and 8). Data set 7 was collected on a Falcon 4i camera (96,000 x magnification, 0.82 Å pixel size). For data sets 9, 10 and 11, the microscope had been upgraded to include a Selectris energy filter in addition to the Falcon 4i camera; movies were recorded at 165,000 x corresponding to a calibrated pixel size of 0.72 Å/pixel, using a slit width of 10 eV. Defocus values ranged from -0.75 µm to -2.25 µm. Movies were recorded in counting mode with a total dose of ∼50-60 e- per Å^2^. Further details of data collection are summarized in Extended Data Table 1

### Cryo-EM data processing

All cryo-EM data processing was performed using the CryoSPARC v3.2 pipeline^35^. Movie stacks were aligned using patch motion correction followed by defocus estimation with patch CTF estimation. For the RT-T/P complex (data set 1), a subset of 200 images was used for automatic particle picking using a blob picker with 60 Å and 130 Å as minimum and maximum particle diameters, respectively. After extensive 2D classification, a selection of the best 2D class averages was used as templates to pick particles from the full data set. Particles were subjected to reference-free 2D classification. A subset of 20 % of particles selected after 2D classification were used for heterogeneous *ab initio* reconstruction with 2 classes. One of the classes resembles the expected 3D map for RT while the other class seems to deviate (Fig. S2). We used these two classes to represent a “good” class and a “junk” class for 3D heterogeneous refinement. After this clean-up, the final “good” particle set was used for non-uniform 3D Refinement^36^. For the remaining data sets, the 3D reconstruction of data set 1 was used to generate projection-based templates for automatic particle picking. After reference-free 2D classification, particles were submitted to 3D heterogenous refinement using the two classes generated *ab initio* for the RT-T/P complex (data set 1). This strategy allowed us to remove “junk” particles still present in the stack after 2D classification by sinking them into the “junk” 3D class. Additional 3D classification with a single reference did not detect underlying heterogeneity or improve the final reconstruction. Cleaned particle stacks were used for non-uniform 3D Refinement. Details of data processing are shown in Figs. S2-S12 and Extended Data Table 1. Resolutions were estimated by Fourier shell correlation using the 0.143-criterion according to the “gold-standard” method. Local resolutions were also calculated in cryoSPARC using BlocRes algorithm^37^ (Extended Data Fig. 1). The reported resolution in the active site was calculated based on the local resolution maps. Fragments of the cryo-EM maps were extracted in Coot v0.9^38^ with a 10 Å cut-off using the α-phosphate of the dATP molecule as the center of the extraction box. The extracted local resolution maps were used to calculate the average local resolution value in the active site using ChimeraX v.1.5^39^ (Extended Data Table 1 and Extended Data Fig. 1).

### Model building and refinement

The X-ray structure of RT-T/P complex (PDB: 5D3G) was used as starting model and docked into the various cryo-EM maps by MOLREP^40^ inside the CCP-EM interface. A series of sharpened maps (B-factors -20, -40, -60, -80, -100, -120 Å^2^) were obtained with the sharpening tool of cryoSPARC and used to manually adjust the model with Coot v0.9^38^. Mg^2+^ ions and dATP were placed into the densities in unsharpened maps. Sharpened maps were used to model side chain density and the models were then refined with PHENIX v1.18.2^41^ using real space refinement. After this refinement, each residue of the model, in particular those of the active site, were manually checked and refined in Coot. Structural models were validated with MolProbity^42^. The final refinement statistics were obtained against unsharpened maps and are summarized in Extended Data Table 1. Electrostatic potentials were calculated using the APBS plugin inside Pymol v.2.5.0 (Schrödinger, LLC).

### Pre-steady-state kinetic analyses

Rapid quench experiments were carried out using a KinTek RQF-3 instrument (KinTek Corporation, Clarence, PA, USA), as described previously^43^. Pre-steady-state data were fitted by nonlinear regression using Sigma Plot software (Jandel Scientific). For the burst experiments, the following equation was used: [T/P+1] = *A*[1 – exp(−*k*1*t*) + *mt*), where *A* represents the burst amplitude, *k*1 the burst rate, and *m* the slope. For single turnover experiments, the following equation was used: [T/P+1] = *A*(1-e^-kobst^). *K_d_* and *k*pol values were calculated by fitting the observed single rate constants (*k*obs) obtained at different concentrations of dNTP to the hyperbolic expression as follows: *k*obs = *k*pol[dNTP]/(*K_d_* + [dNTP]), where *K_d_* is the equilibrium dissociation constant for the interaction of dNTP with the RT-T/P complex and *k*pol is the maximum first order rate constant for dNTP incorporation.

## Data availability

Cryo-EM maps and coordinates have been deposited in the Electron Microscopy Data Bank (EMDB) and the Protein Data Bank (PDB), respectively, with accession numbers EMD-43114 and 8VB6 (RT-T/P); EMD-43115 and 8VB7 (RT-T/P-DNA state R-1); EMD-43116 and 8VB8 (RT- T/P-DNA state R-2); EMD-43117 and 8VB9 (RT-T/P-DNA state R-2’); EMD-43120 and 8VBC (RT-T/P-DNA state T8sec); EMD-43121 and 8VBD (RT-T/P-DNA T’8sec); EMD-43122 and 8VBE (RT-T/P-DNA P2min); EMD-43123 and 8VBF (RT-T/P-DNA P6min); EMD-43124 and 8VBG (RT- T/P-dApCpp); EMD-43125 and 8VBH (RT-T/P-DNA No MgCl2); and EMD-43126 and 8VBI (RT-T/P-DNA 3’-terminated primer)

## Supporting information

Cryo-EM processing workflow

## Acknowledgments

We thank Dr. Craig D. Kaplan for his critical reading of the manuscript. This research was supported by NIH Grants P50 GM082251 and R01 AI175067 (to G.C.). All Cryo-EM data were collected at the Cryo-Electron microscopy Facilities at the Department of Structural Biology. The University of Pittsburgh Titan Krios microscope and Falcon 3 camera were supported by the Office of the Director, National Institutes of Health, under award numbers S10 OD025009 and S10 OD019995. The content is solely the responsibility of the authors and does not necessarily represent the official views of the National Institutes of Health.

## Author contributions

X.Z. performed protein purification. S.V. prepared cryo-EM samples. S.V. and J.F.C. collected and processed cryo-EM data. S.V. and G.C. performed model building. U.S. contributed to structure analysis. N.S.C. performed pre-steady-state kinetic analyses. S.V., N.S.C. and G.C. wrote the manuscript with input from all co-authors. G.C. conceived the project and together with N.S.C. coordinated and supervised experimental work and data analysis.

---

**Extended Data Fig. 1.**
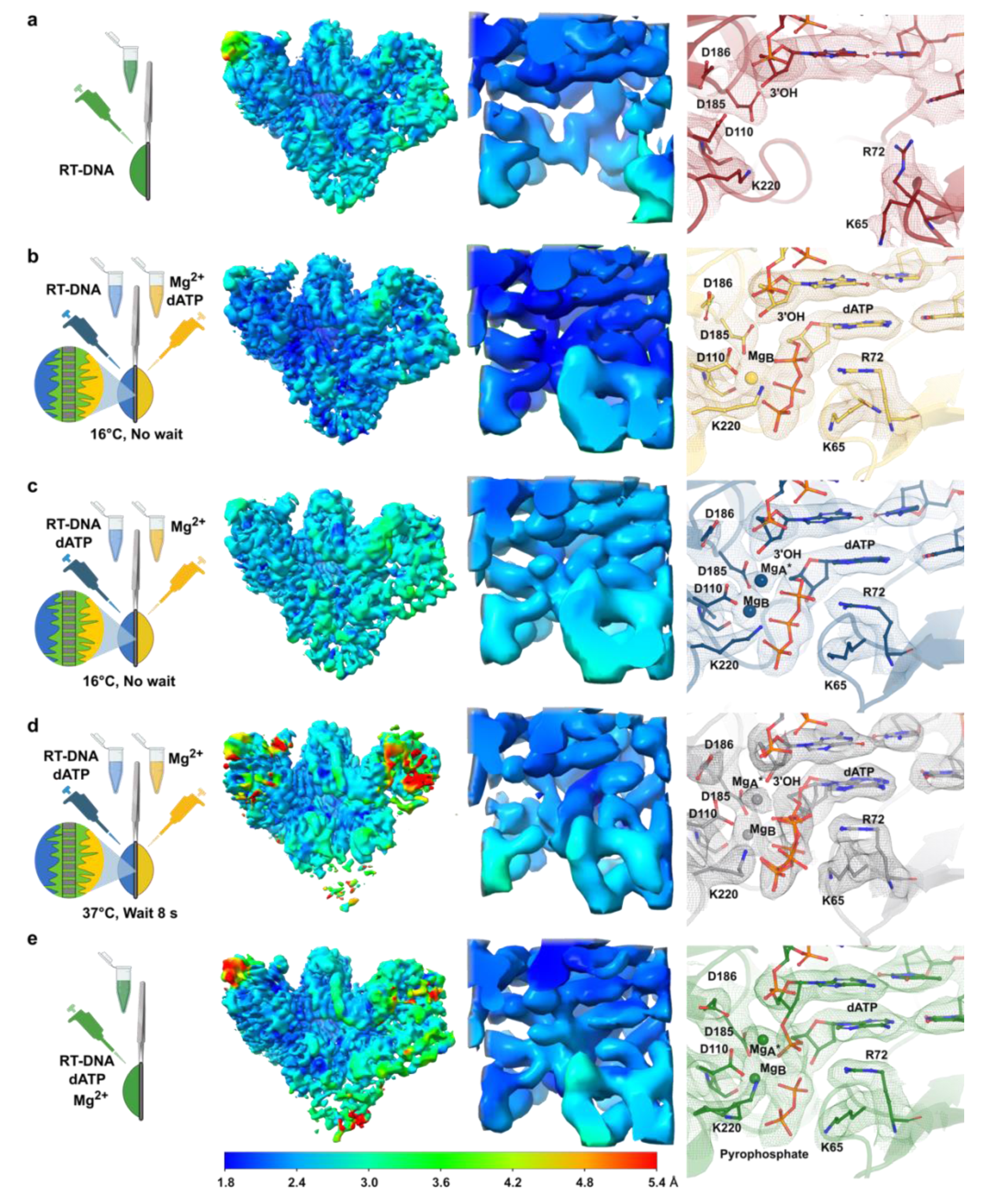
Overall quality of Cryo-EM maps. **a**, dNTP-free RT-T/P complex and states (**b**) R-1, (**c**) R-2, (**d**) T8sec and (**e**) P6min of the RT-T/P- dATP complex. From left to right: first column, schematic representation of the grid preparation protocol with R-1, R-2, and T8sec obtained with *in situ* “diffusion/blot” experiments. P6min and P2min (not shown) were obtained by incubating all components of the reaction at 37 °C for 6 min or 2 min prior to solution application on grid. Second column, surface of cryo-EM maps colored according to local resolution; third column, local resolution in the active site; fourth column, model in the active site fit into the density map (mesh representation).

**Extended Data Fig. 2:**
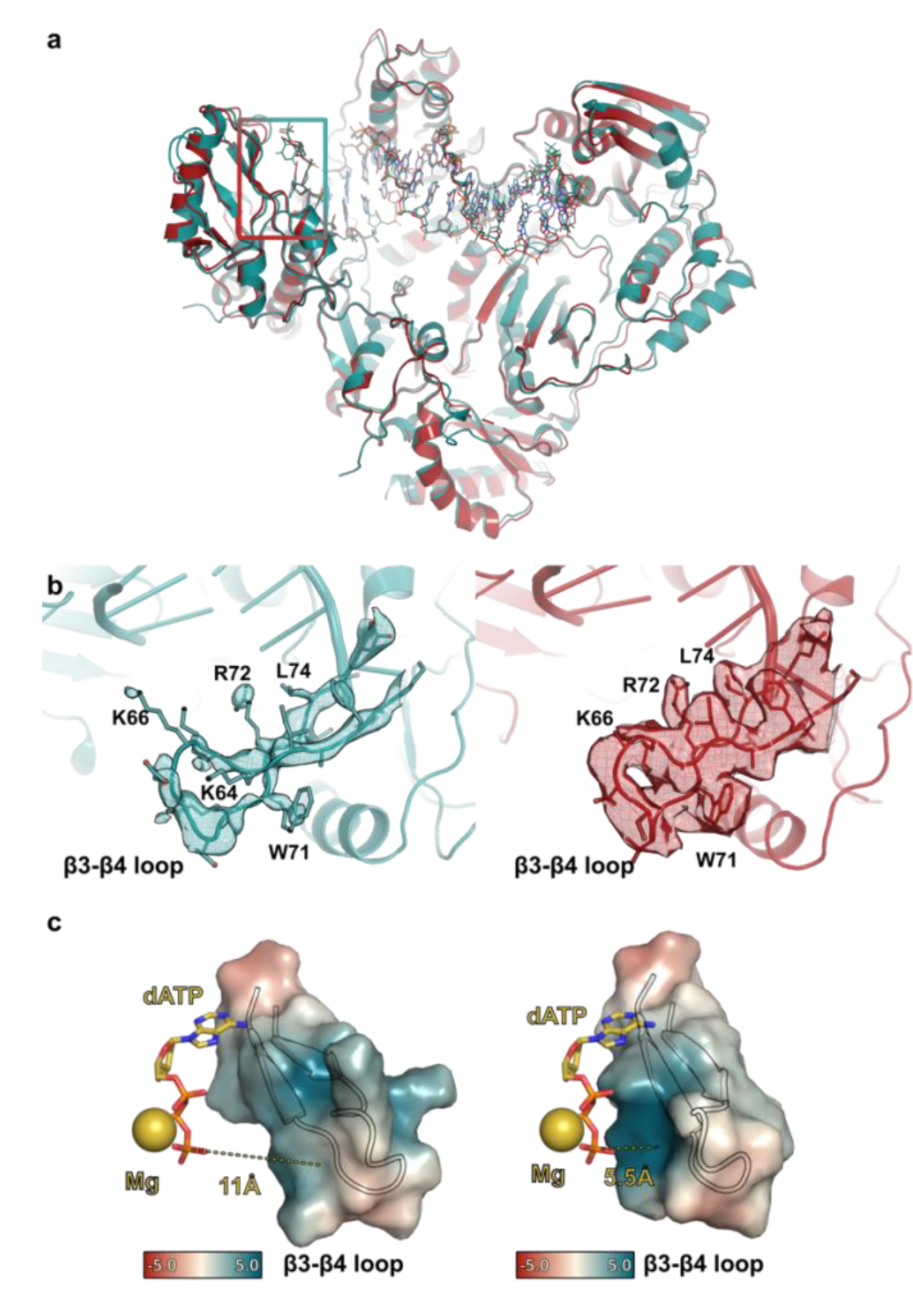
Comparison of HIV-1 RT structure by Cryo-EM and X-ray. **a**, Cartoon representation of the cryo-EM structure of RT-T/P binary complex (red) superimposed to the corresponding X-ray structure with same DNA aptamer (PDB: 5D3G, teal). DNA aptamer displayed in stick representation. Square indicates active site. **b**, Comparison of density map for the β3-β4 loop observed in X-ray (teal, map countered at σ = 0.9) or in cryo-EM (red, map contoured at σ = 7). Protein backbone is shown in cartoon representation with relevant residues displayed in stick. Cryo-EM map allows modeling of all important residues in the β3-β4 loop. **c**, Surface representation of the β3-β4 loop in the RT-T/P complex (left) or the state R-1 of the RT- T/P-dATP complex (right) colored by electrostatic surface potential (from red -5 kT/e to blue +5 kT/e), showing the positively charged region that could interact with the negatively charged phosphate tail. dATP and Mg^2+^ displayed in ball-and-stick representation as model in state R-1.

**Extended Data Fig. 3:**
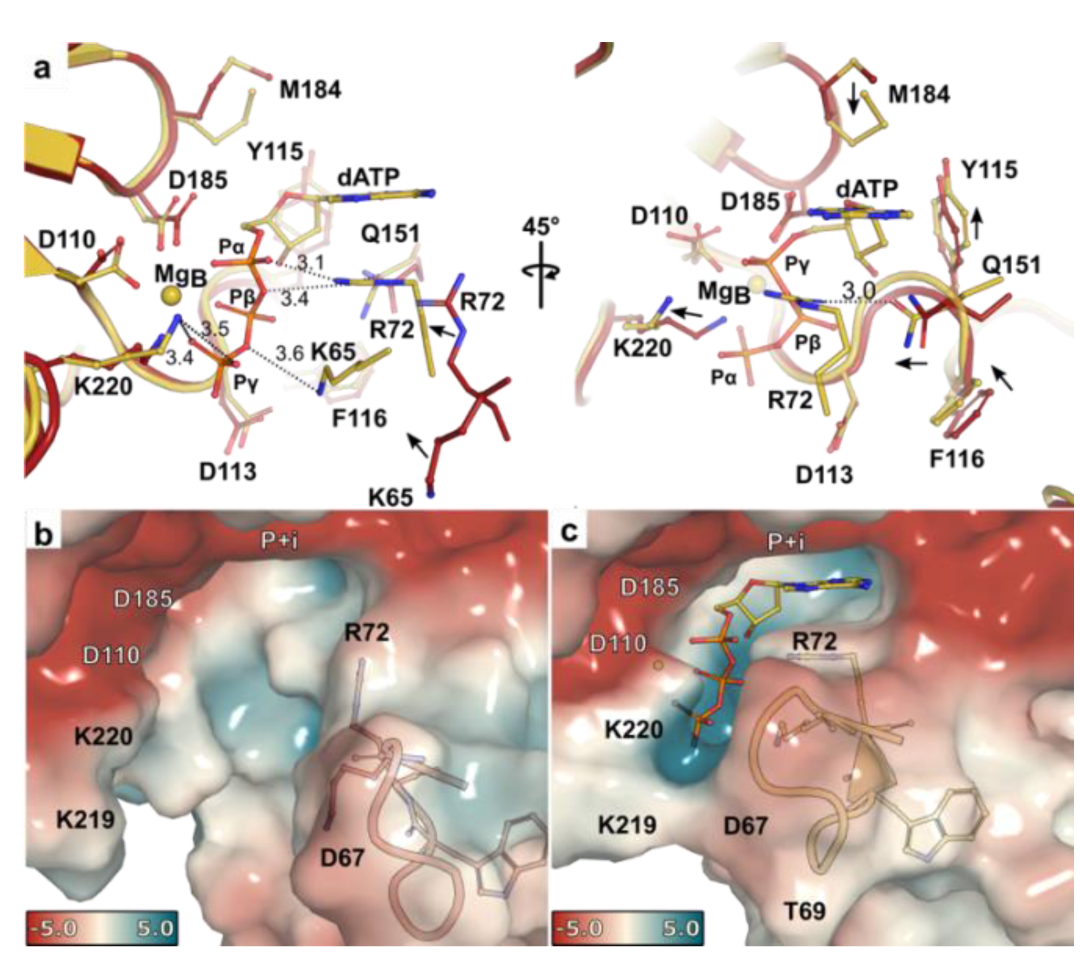
Comparison of dATP-free RT-T/P complex and dATP-bound RT-T/P complex. **a**, Two views of the overlay between the RT-T/P complex (red) and state R-1 of the RT-T/P-dATP complex (yellow). The incoming dATP is stabilized by salt bridges between the phosphate tail and residues K220, K65 and R72 (left); additional conformational changes of key residues in the dNTP-binding pocket include M184, Q151, Y115, F116 (right). Coordination distances (in Å) are indicated next to dashed lines. **b,c**, Surface representation of (b) the RT-T/P complex and **(c)** the state R-1 of the RT-T/P-dATP complex colored according to electrostatic potential from red (-5 kT/e, negative charge) to blue (+5 kT/e, positive charge). Electrostatic potentials were calculated in the absence of dATP. β3-β4 loop displayed in cartoon with representative residues shown in ball-and-stick. A perfectly fitting “blue” crevice is formed around the contour of dATP.

**Extended Data Fig. 4:**
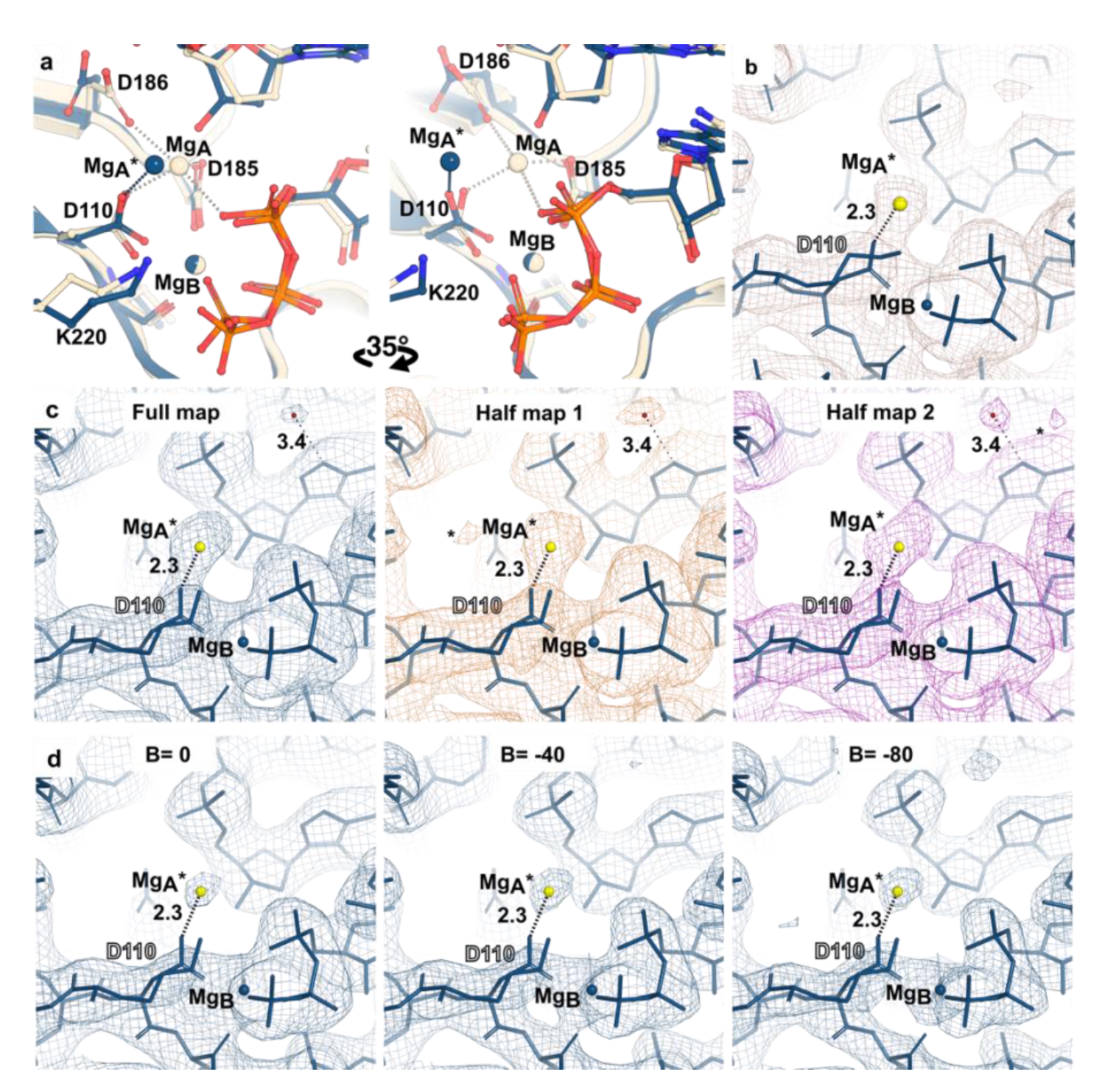
Cryo-EM observation of metal ion A* in RT-T/P dATP complex. **a**, Active site superposition of state R-2 of RT-T/P-dATP complex (blue) with an X-ray structure of HIV-1 RT-RNA/DNA ternary complex with dATP (PDB: 4PQU, cream). The X-ray model of RT has a chain-terminating primer (no 3’-OH). The position of Mg^2+^ ion in site A* does not coincide with the Mg^2+^ ion site A observed in the X-ray structure. Coordination of Mg^2+^ is marked with dashed lines (distances lower than 2.6 Å). **b**, Position of Mg^2+^ ion in site A* was reproduced in a second data set with same experimental conditions for state R-2 of RT-T/P-dATP (R-2’ or data set 4 in Extended Data Table 1). Unsharpened map (light brown mesh) displayed at a threshold of 11 σ. **c**, Full map (unsharpened) and corresponding half-maps of state R-2. The displayed model was refined against full map. Strong features have common intensity and position in both half- maps whereas background noise (*) can be observed in one half-map. Density that may correspond to waters (red dot) also displayed high mobility between half-maps. Maps are displayed at a threshold of ∼9 σ. **d**, Full map displayed at various values of B-factor (Å^2^) sharpening. Maps are displayed at a threshold of ∼14 σ. The position and size of the density for Mg^2+^ ion A* remains similar for all sharpening values. Mg^2+^ in ion site A* is colored in yellow in **b-d**. Coordination distances (in Å) are indicated next to dashed lines.

**Extended Data Fig. 5:**
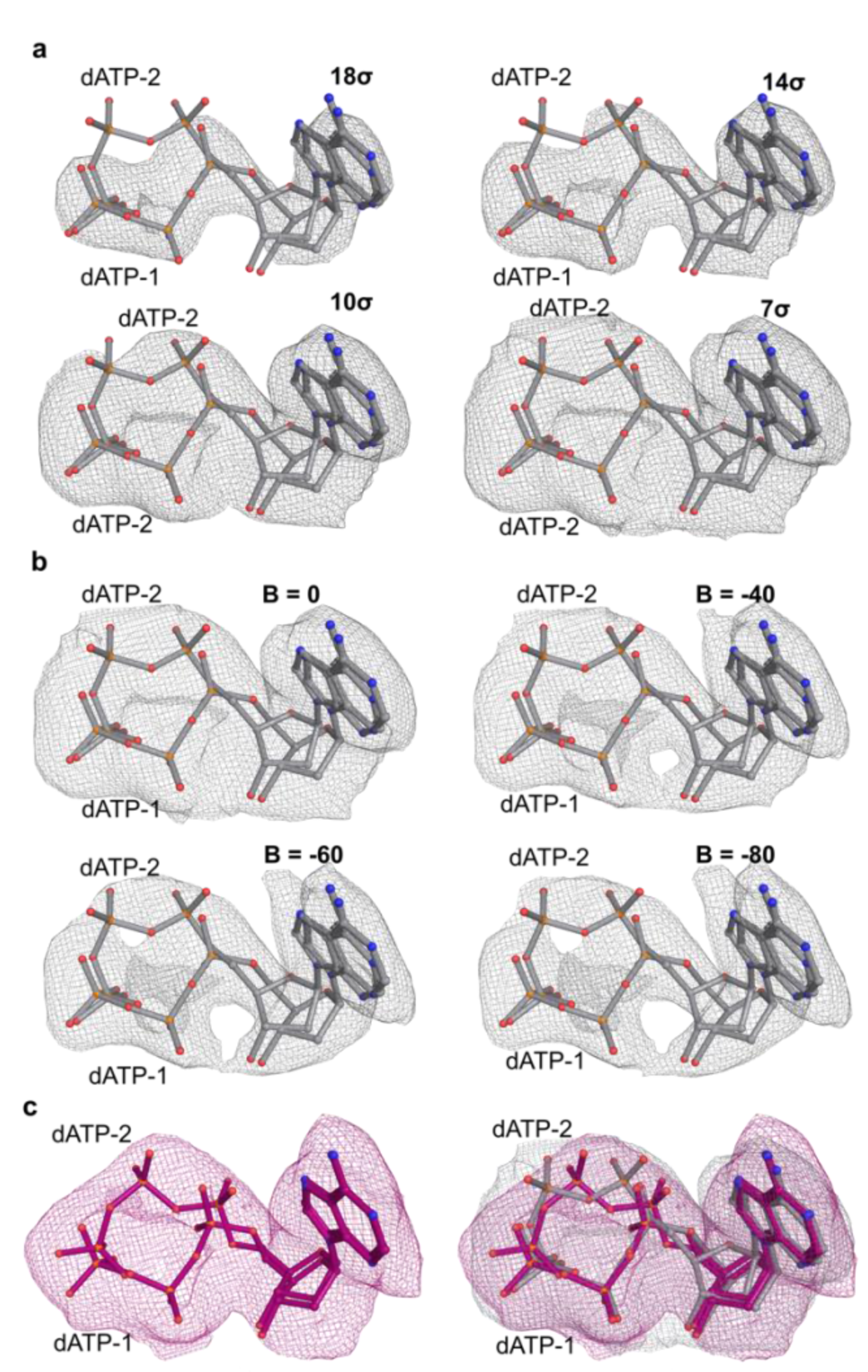
Two conformations of phosphate tail in the transition state of the RT-T/P-dATP complex. **a**, Cryo-EM density map (gray mesh, unsharpened) displayed at several thresholds in state T8sec. Extra density emerges at lower thresholds that can only be explained by two conformations of the phosphate tail (shown in ball-and-stick). Density observed for dATP-1 state at high sigma values of the map correlates with a higher occupancy refinement in Phenix (0.7 for dATP-1 vs 0.3 for dATP-2). **b,** Cryo-EM density map of T8sec displayed at different B-factor (Å^2^) sharpenings (contoured at ∼7 σ). **c,** Left, state T’8sec of the RT-T/P-dATP (after 8 seconds of incubation at 30 °C on the grid). Right, overlay of dATP structure model in T’8sec (purple) and T8sec (gray). Maps are displayed both at threshold of 7.2 σ and unsharpened.

**Extended Data Fig. 6.**
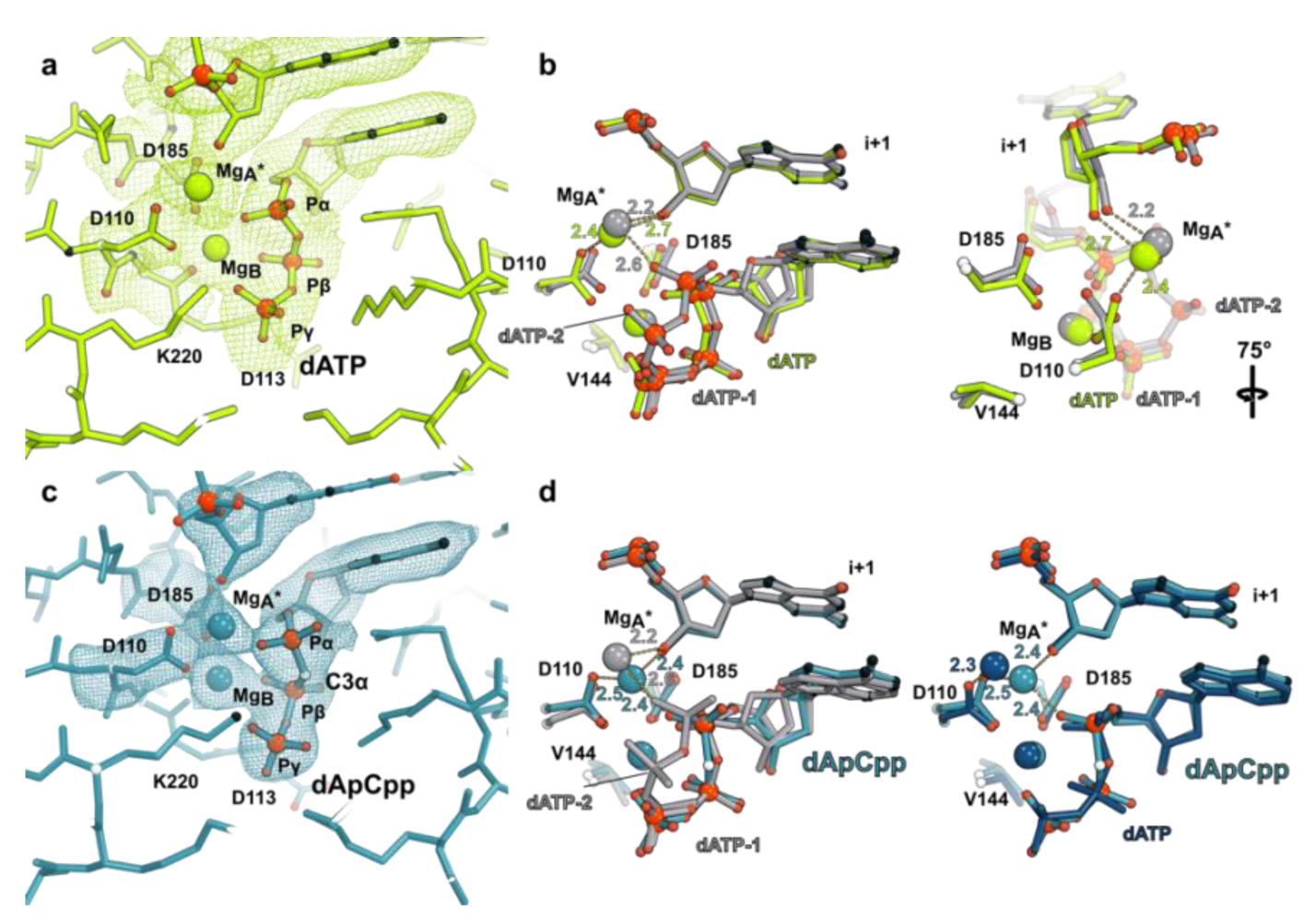
Structures of the early product state (P2min) and dApCpp. **a**, Cryo-EM density map of the state P2min of the RT-T/P-dATP complex (lime green mesh) with corresponding model in ball-and-stick. A strong density for Mg^2+^ in sites B and A* can be observed. Map displayed with sharpening of B-factor = -20 Å^2^ at contoured level of 5 σ. **b**, Overlay of active sites of states P2min (lime green) and T8sec (gray). **c**, Density map of the active site of RT- T/P bound to a non-hydrolyzable dNTP (dApCpp) displayed unsharpened and contoured at 10 σ (teal mesh). **d**, Comparison of metal positions in RT-T/P-dApCpp (teal) and RT-T/P-dATP complexes. State T8sec (gray), left; state R-2 (blue), right. Coordination distances (in Å) are indicated next to dashed lines.

**Extended Data Fig. 7.**
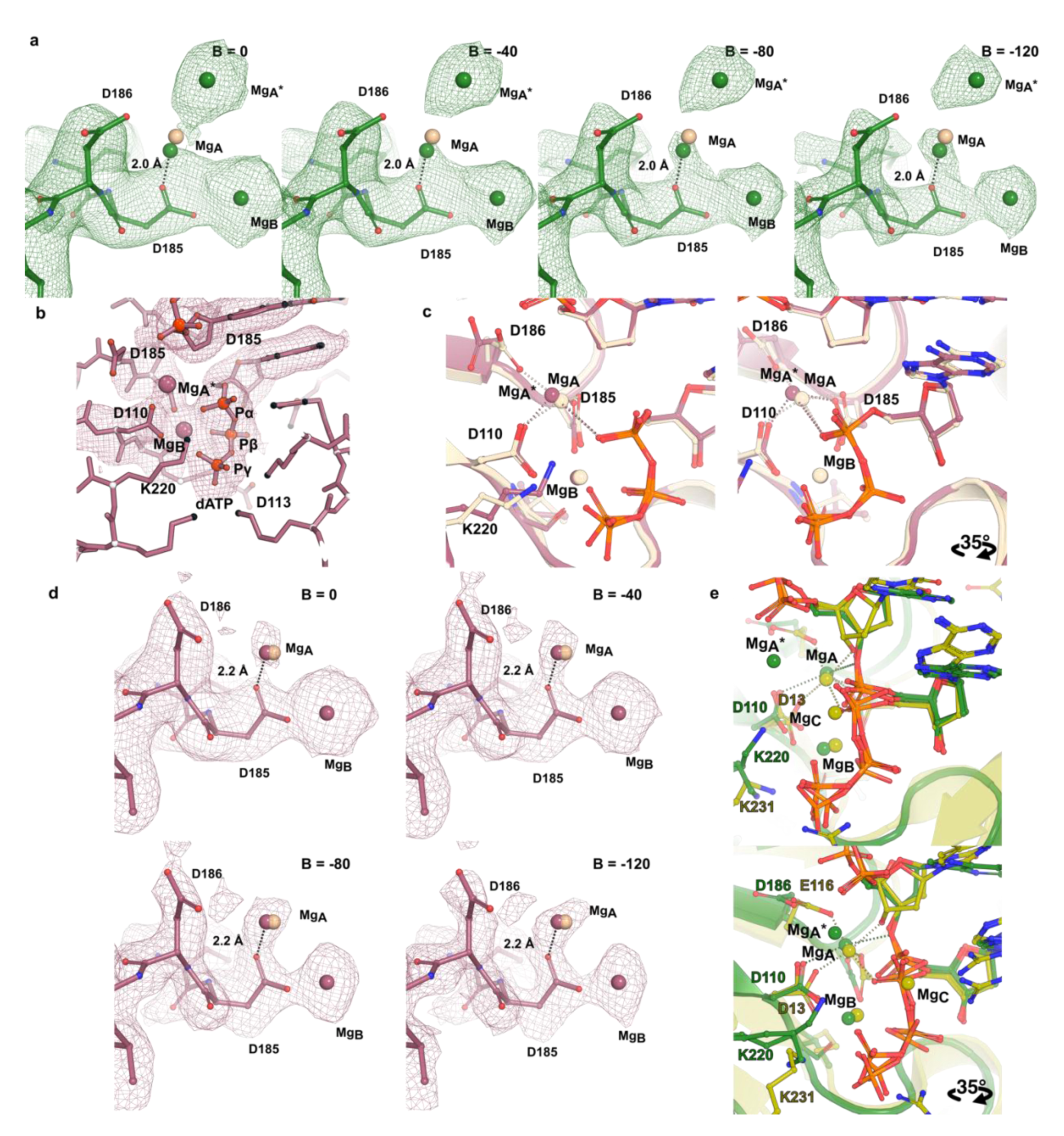
Observation of density for site A Mg^2+^. **a**, Density for metal A Mg^2+^ site, in addition to sites A* and B, can be observed in state P6min if high sharpening is applied. Maps are displayed at a threshold of ∼5.5 σ. Sphere in cream represents the position for Mg^2+^ site A reported in the X-ray structure of RT-T/P-dNTP (PDB: 4PQU, 3’ chain-terminated). **b**, Cryo-EM density map of the active site on RT-T/P-dATP-Mg^2+^ in the absence of 3’-OH. Density is detected for site A Mg^2+^ but not for site A*. Unsharpened map contoured at 6.5 σ. **c**, Active site superposition of the cryo-EM structure obtained with a 3’ chain-terminated primer (violet) with the corresponding X-ray structure (PDB: 4PQU, cream) showing that the position of site A Mg^2+^ is similar for both structures. **d**, Density for site A metal in the cryo-EM structure of RT-T/P-dATP in the presence of 3’ chain terminated primer at different values of map sharpening. Maps are displayed at ∼7 σ. **e**, Overlay of state P6min in RT-T/P-dATP complex (green) with Human DNA polymerase η, pol η-DNA-dATP ternary complex (PDB: 4ECX, Lime). Positions of Mg^2+^ at sites A and B are similar for both polymerases but pol η active site features a third Mg^2+^ site C that is not observed in RT. Coordination of Mg^2+^ is marked with dashed lines for distances smaller than 2.6 Å in (**c**) and (**d**).

**Extended Data Fig. 8:**
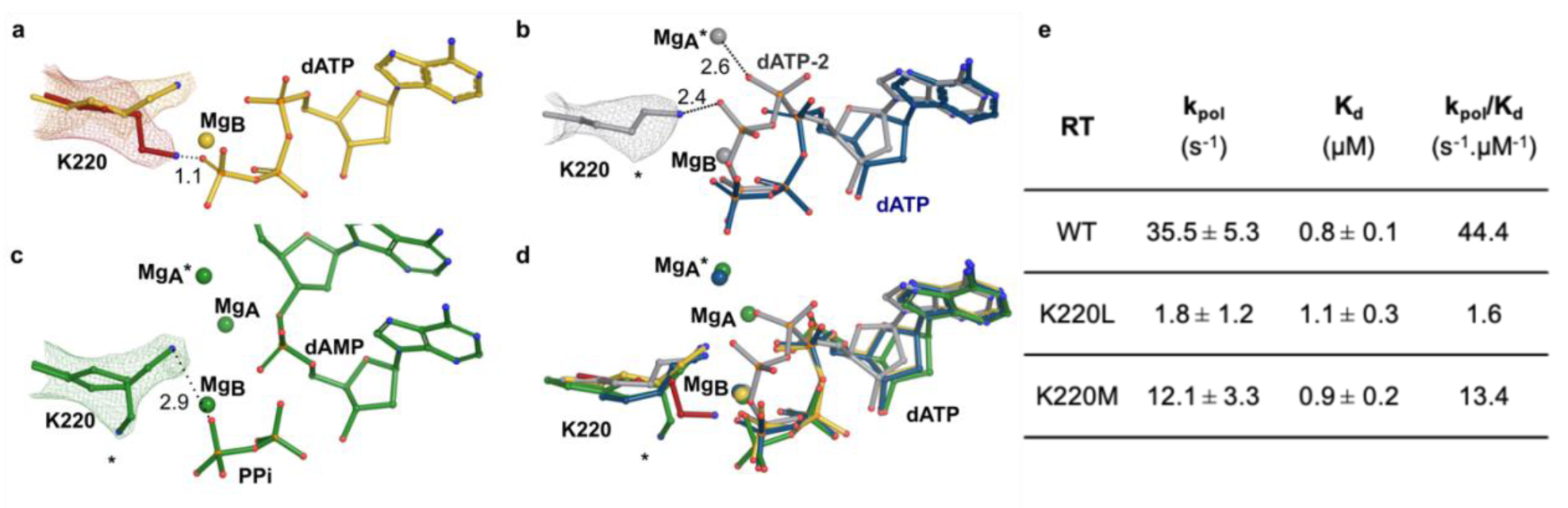
Positioning and interaction of K220 with substrates and products in the RT active site in the different RT-T/P-dATP complexes. Structural changes of K220 during HIV-1 RT reaction. **a**, The arrival of dATP to the active site (yellow, state R-1) forces a repositioning in K220 (red, dATP-free state) to avoid clashes. **b**, K220 directly interacts with the phosphate tail of dATP-2 prior to nucleotide addition (blue, state R-2; gray, state T8sec). **c**, Following translocation (green, state P6min), a second conformation for K220 is detected (marked with *). **d**, Overlay of all stages showing the movement of K220 during dATP addition cycle. **e**, Transient kinetic analysis of nucleotidyl transfer by wild-type (WT), K220L and K220M HIV-1 RT. Coordination distances (in Å) are indicated next to dashed lines

**Extended Data Fig. 9:**
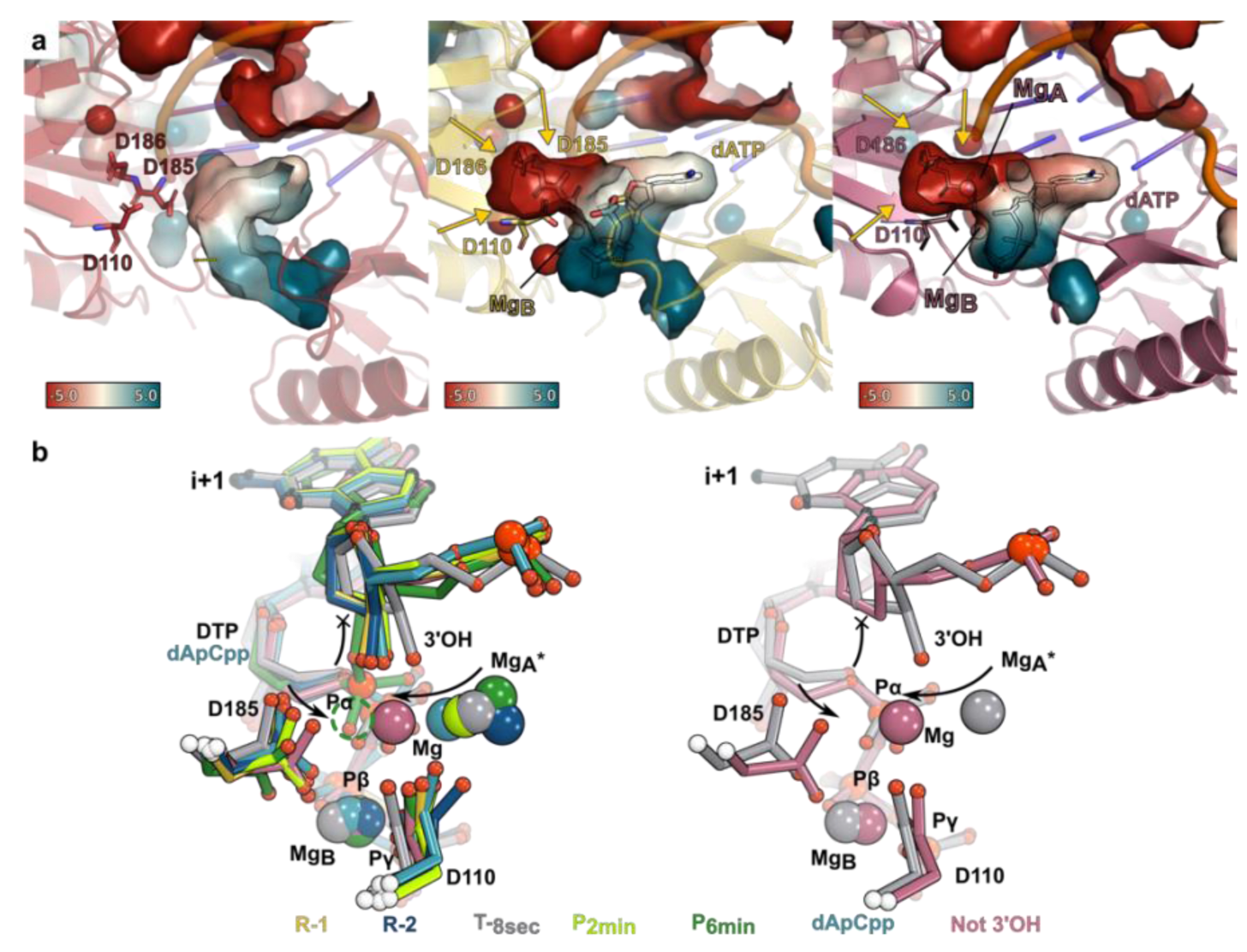
High mobility of Mg^+2^ at site A*. **a,** Electrostatic surface potential of the dNTP binding site. Graphical surface representation of the charged cavities colored by electrostatic potential for RT-T/P binary complex (left), state R-1 of RT-T/P-dATP (center) and RT-T/P-dATP 3’ chain-terminated (right). A negatively charged pocket is formed upon dATP binding (yellow arrows). Pockets were detected using a 4 Å cavity detection radius and a 4 Å solvent radius cavity detection cutoff in Pymol. Protein backbone is shown in cartoon representation with key residues displayed in ball-and-stick representation. **b**, Superposition of all states (left panel) reported in this study showing that the site A* Mg^2+^ has high mobility, while the position of site B Mg^2+^ had minor changes in all experiments. Right panel, overlay of the chain terminated structure (violet) with the T8sec structure shows that the Mg^2+^ ion shifts from the A* to the canonical A position (curved arrow) in the absence of a 3’OH group, with concomitant D110 and D185 residue rearrangement.

**Extended Data Table 1.**
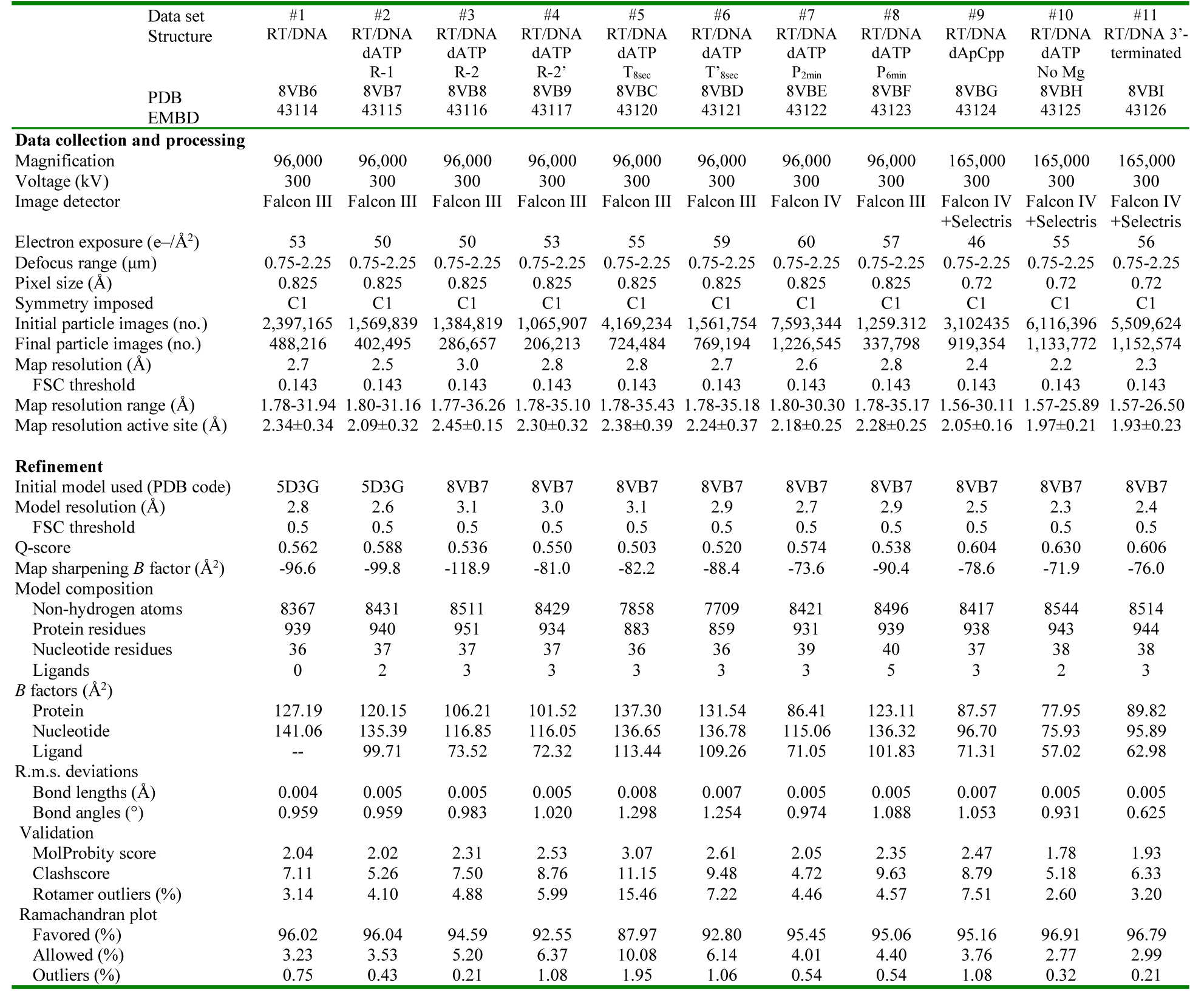
Cryo-EM data collection, refinement and validation statistics.

