## Supplementary material for "Structures of kinetic intermediate states of HIV-1 reverse transcriptase DNA synthesis": Cryo-EM processing workflow

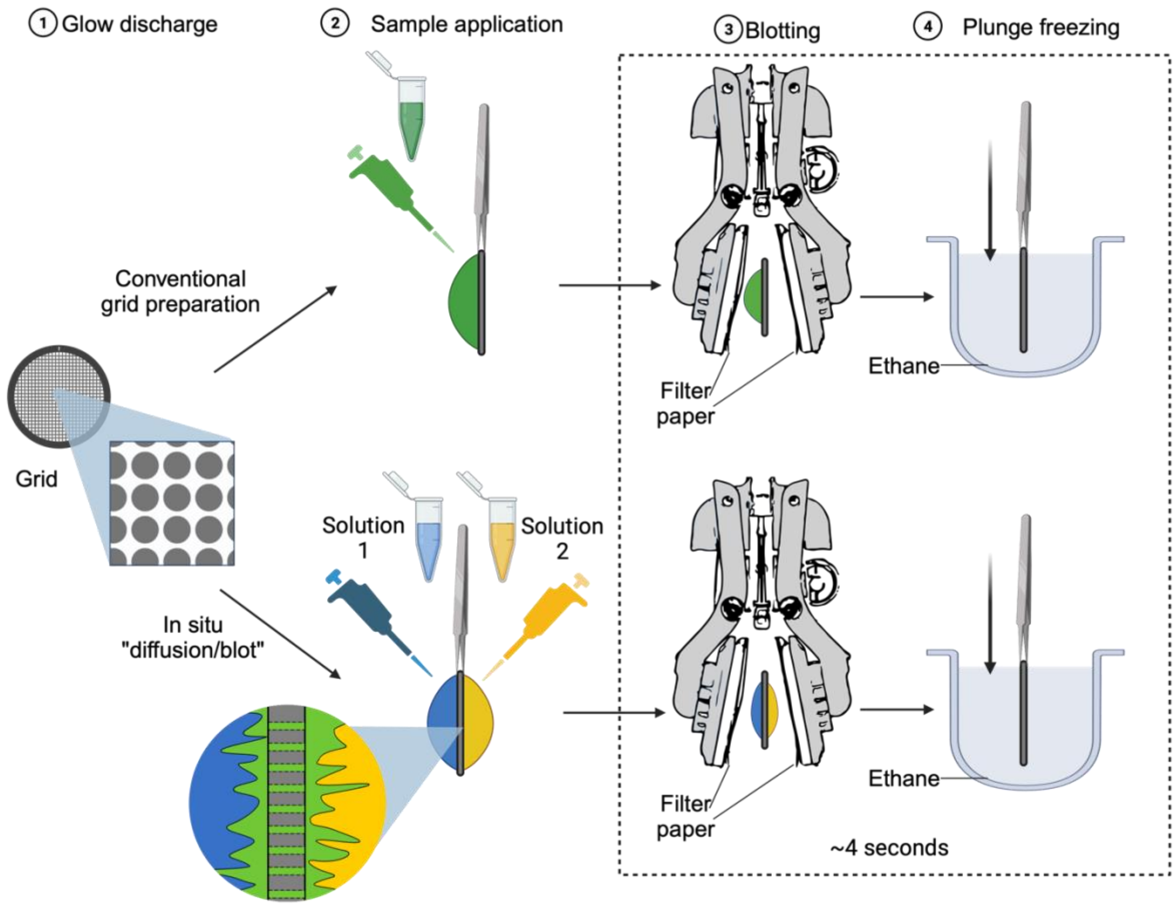


**Fig. S1.** Schematic representation of the cryo-EM grid preparation protocol. Two different protocols were implemented: conventional grid preparation and in situ “diffusion/blot”. In the first one, a stable solution preincubated with all components of the reaction is applied to the grid (Data sets 1, and 7 through 11 in Extended Data Table 1). On the second one, the in situ “diffusion/blot”, two separate solutions (blue and yellow in the scheme) are applied on the opposite sides of the cryo-EM grid and mixing occurs on the grid by diffusion through the matrix of holes (generating green solution in situ). Application of the second solution starts the RT reaction allowing for short reaction times in the scale of seconds (Data sets 2 through 6). The subsequent blotting and plunge freeze cycle takes ~4 seconds.

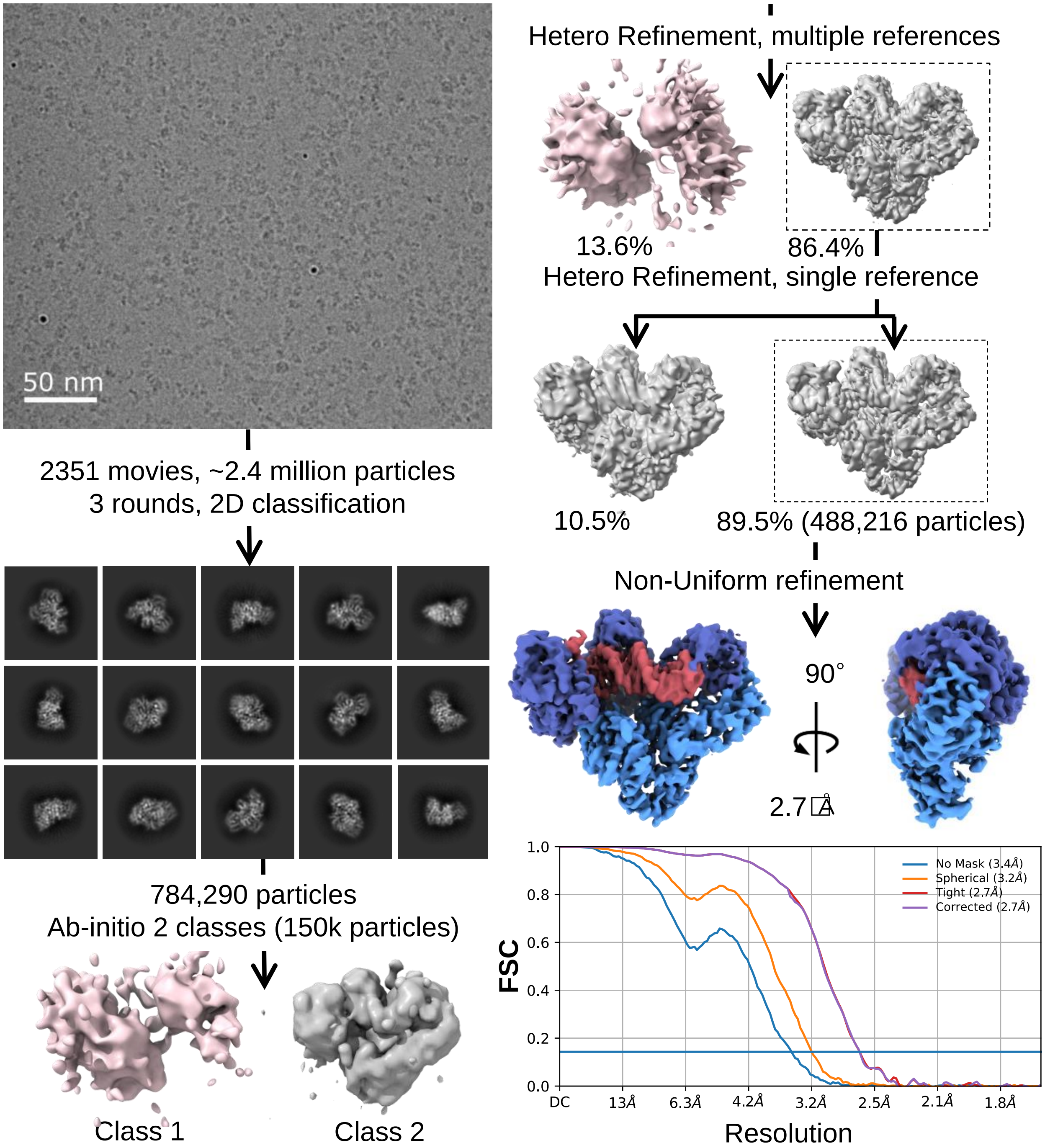


**Fig. S2.** Workflow of Cryo-EM processing of RT-DNA without dATP (Data set 1). Upper-left corner shows a typical micrograph for HIV-1 RT experiments reported in the present study. After 2D classification, a subset of 150,000 particles were used for *ab-initio* 3D reconstruction with two classes. These two classes were later filtered and used as references for 3D classification (For data sets 1 through 8). After 3D classification (heterogeneous refinement), the resulting clean particle stack was used for non-Uniform 3D refinement. All processing was done inside cryoSPARC pipeline. The resolution reported is based on the gold standard Fourier shell correlation (FSC) curve with cut-off value of 0.143.

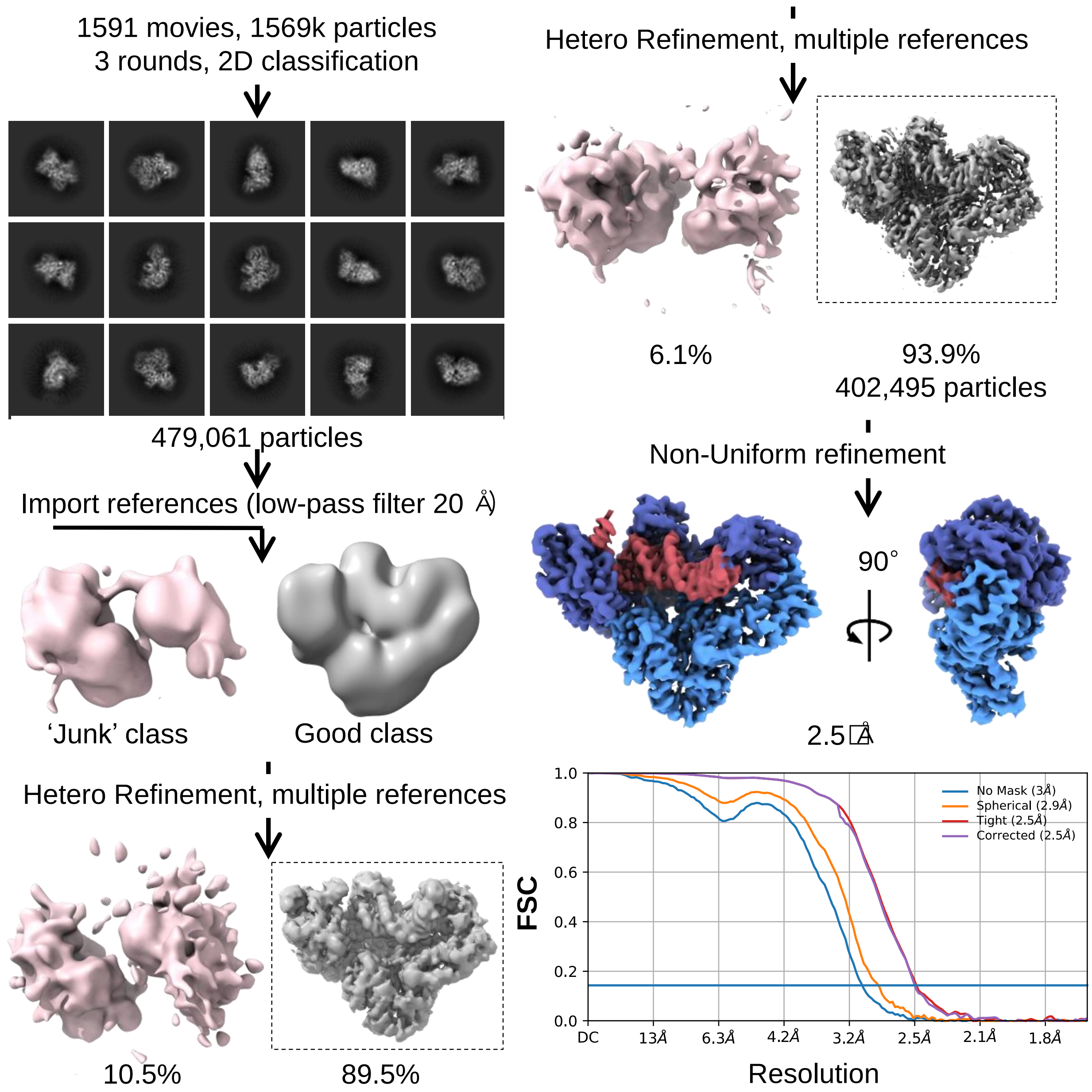


**Fig. S3.** Workflow of Cryo-EM processing of RT-DNA-dATP, State R-1 (Data set 2). After 2D classification and 3D classification (heterogeneous refinement), the clean particle stack is used for non-Uniform 3D refinement. All processing was done inside cryoSPARC pipeline. The resolution reported is based on the gold standard Fourier shell correlation (FSC) curve with cut-off value of 0.143.

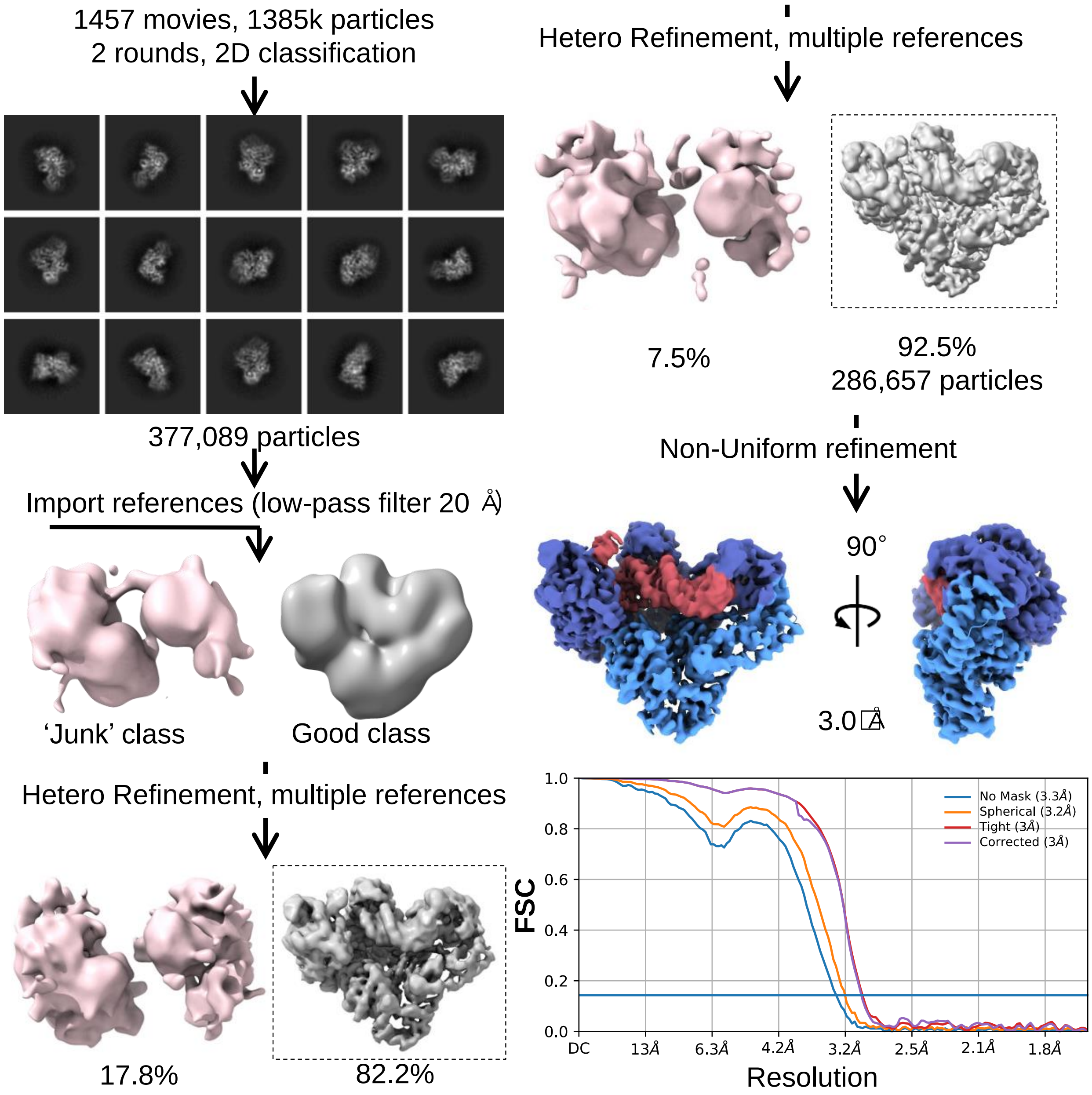


**Fig. S4.** Workflow of Cryo-EM processing of RT-DNA-dATP, State R-2 (Data set 3). After 2D classification and 3D classification (heterogeneous refinement), the clean particle stack is used for non-Uniform 3D refinement. All processing was done inside cryoSPARC pipeline. The resolution reported is based on the gold standard Fourier shell correlation (FSC) curve with cut-off value of 0.143.

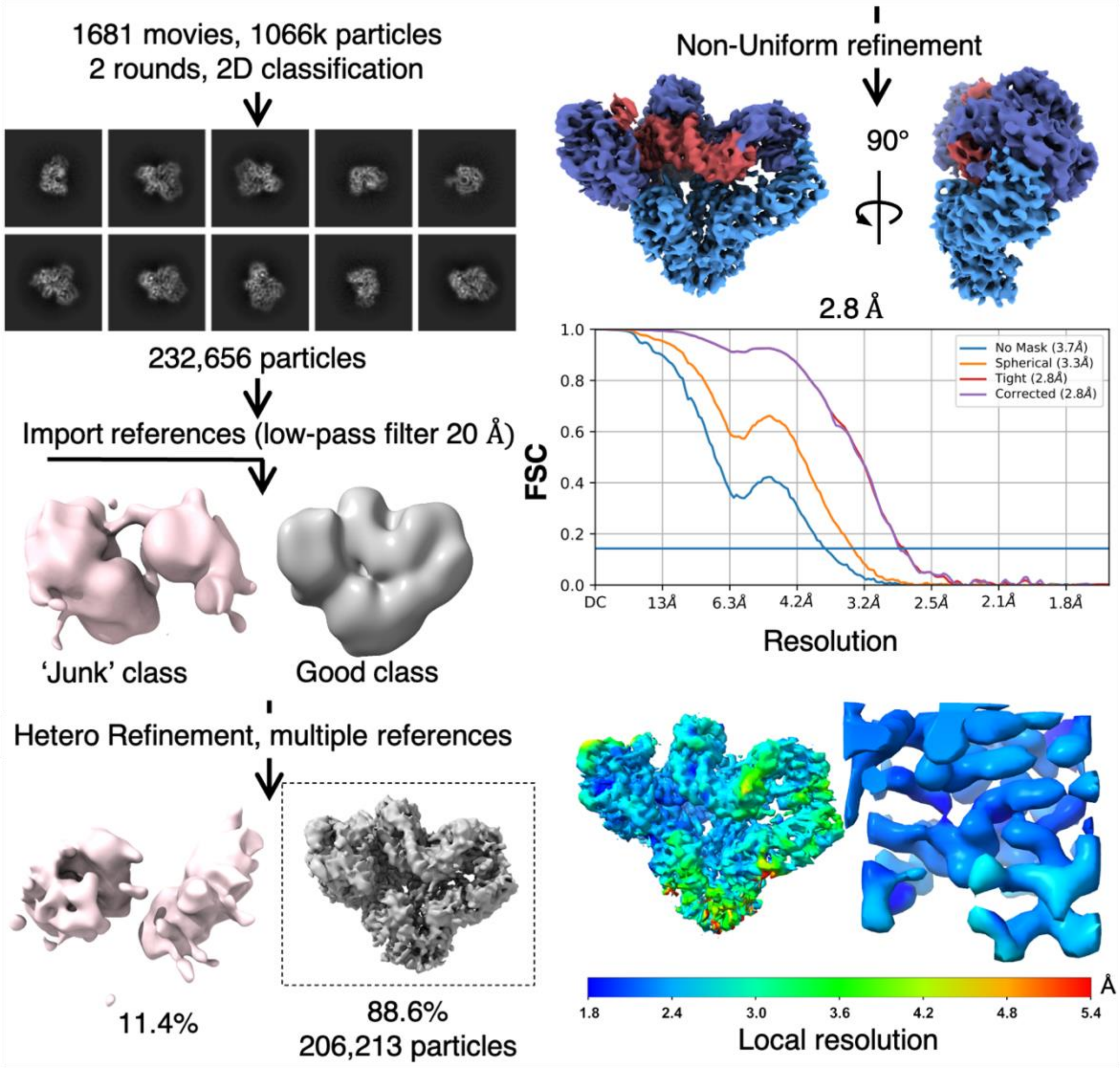


**Fig. S5.** Workflow of Cryo-EM processing of RT-DNA-dATP, State R-2' (Data set 4). Data set 4 implements same experimental condition as Data set 3. After 2D classification and 3D classification, the clean particle stack is used for non-Uniform 3D refinement. All processing was done inside cryoSPARC pipeline. The resolution reported is based on the gold standard Fourier shell correlation (FSC) curve with cut-off value of 0.143.

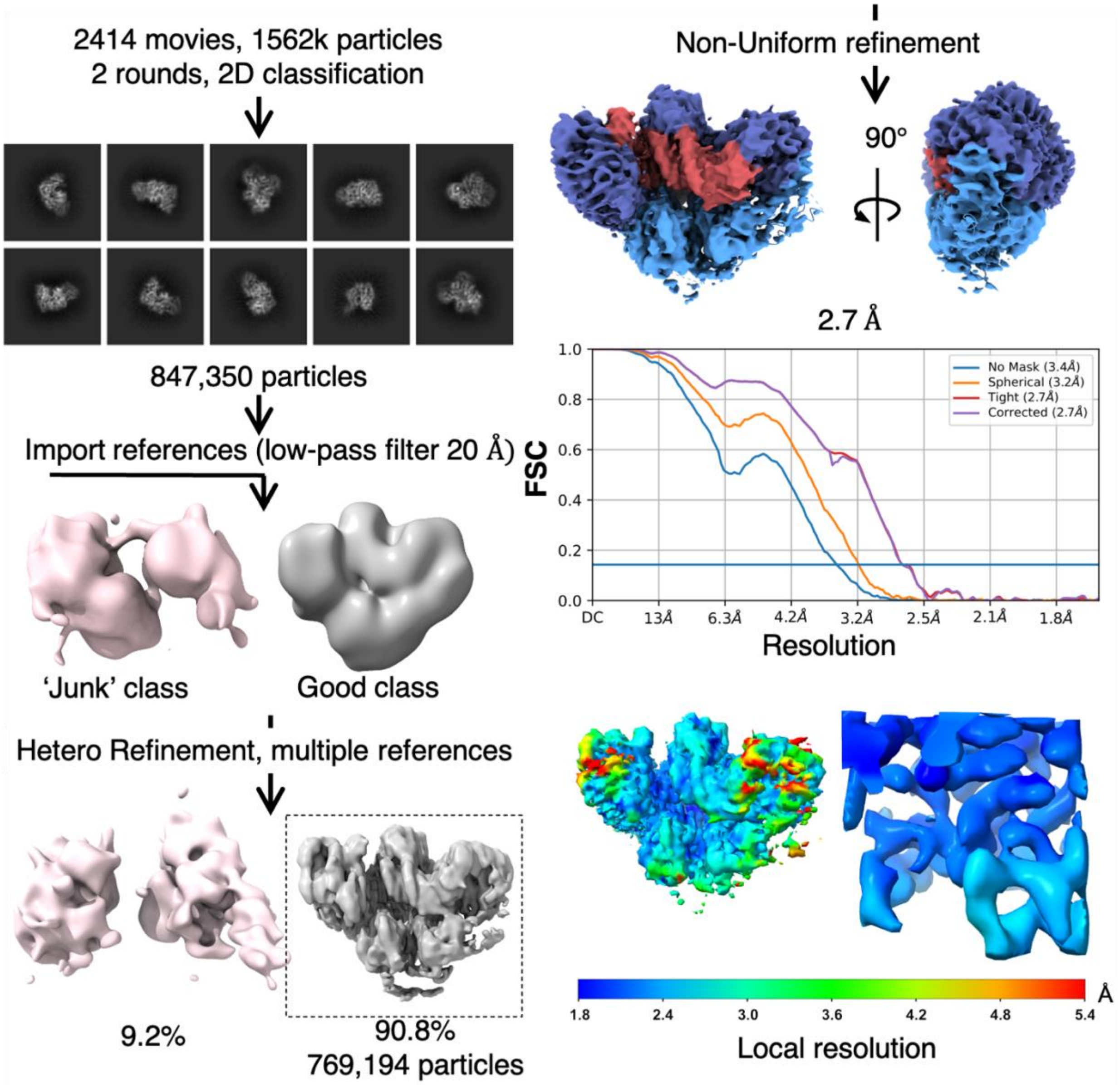


**Fig. S6.** Workflow of Cryo-EM processing of RT-DNA-dATP, State $T_{6sec}$ (Data set 5). After 2D classification and 3D classification (Heterogeneous refinement), the clean particle stack is used for non-Uniform 3D refinement. All processing was done inside cryoSPARC pipeline. The resolution reported is based on the gold standard Fourier shell correlation (FSC) curve with cut-off value of 0.143.

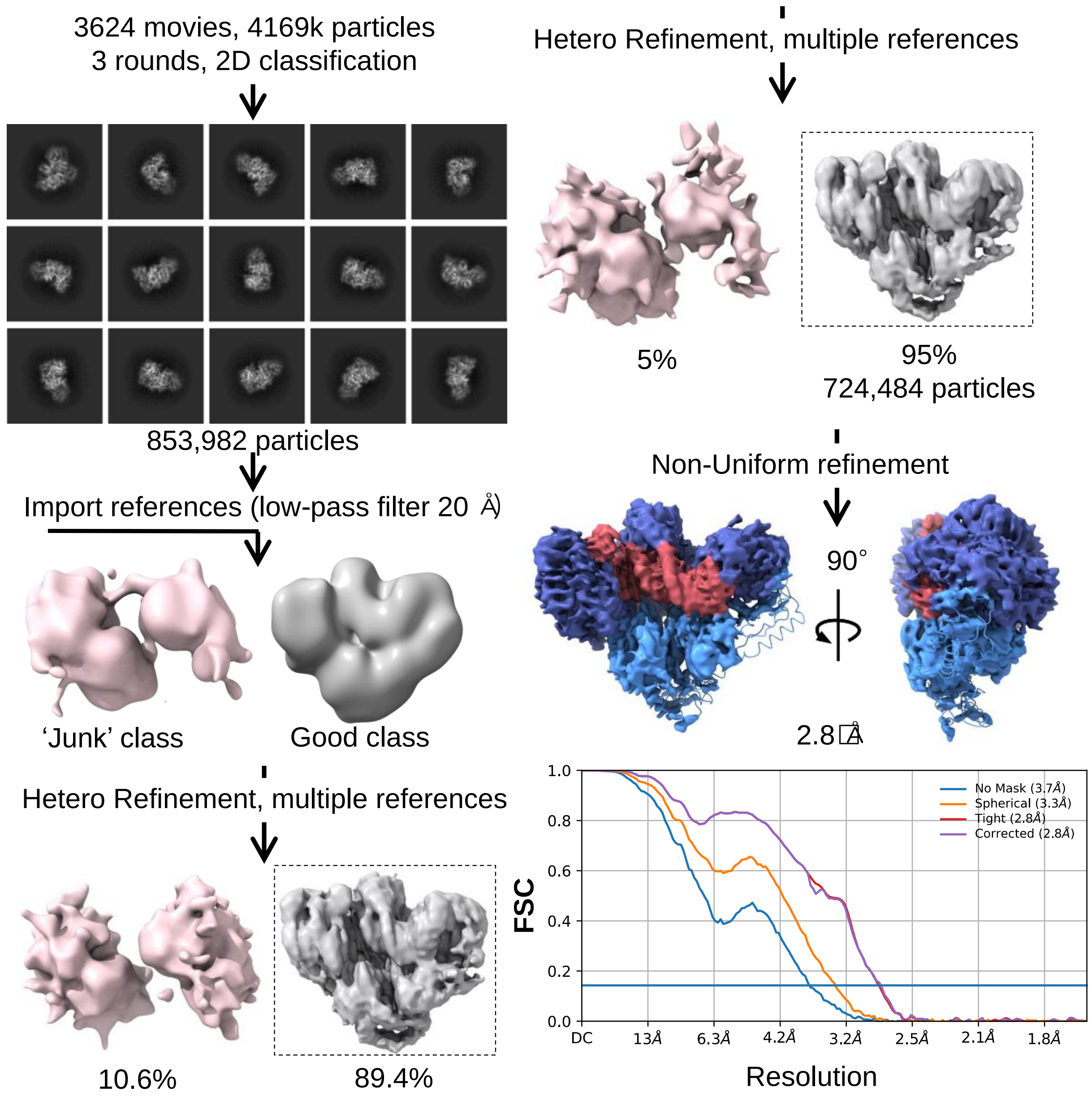


**Fig. S7.** Workflow of Cryo-EM processing of RT-DNA-dATP, State $T_{8sec}$ (Data set 6). After 2D classification and 3D classification (Heterogeneous refinement), the clean particle stack is used for non-Uniform 3D refinement. All processing was done inside cryoSPARC pipeline. The resolution reported is based on the gold standard Fourier shell correlation (FSC) curve with cut-off value of 0.143.

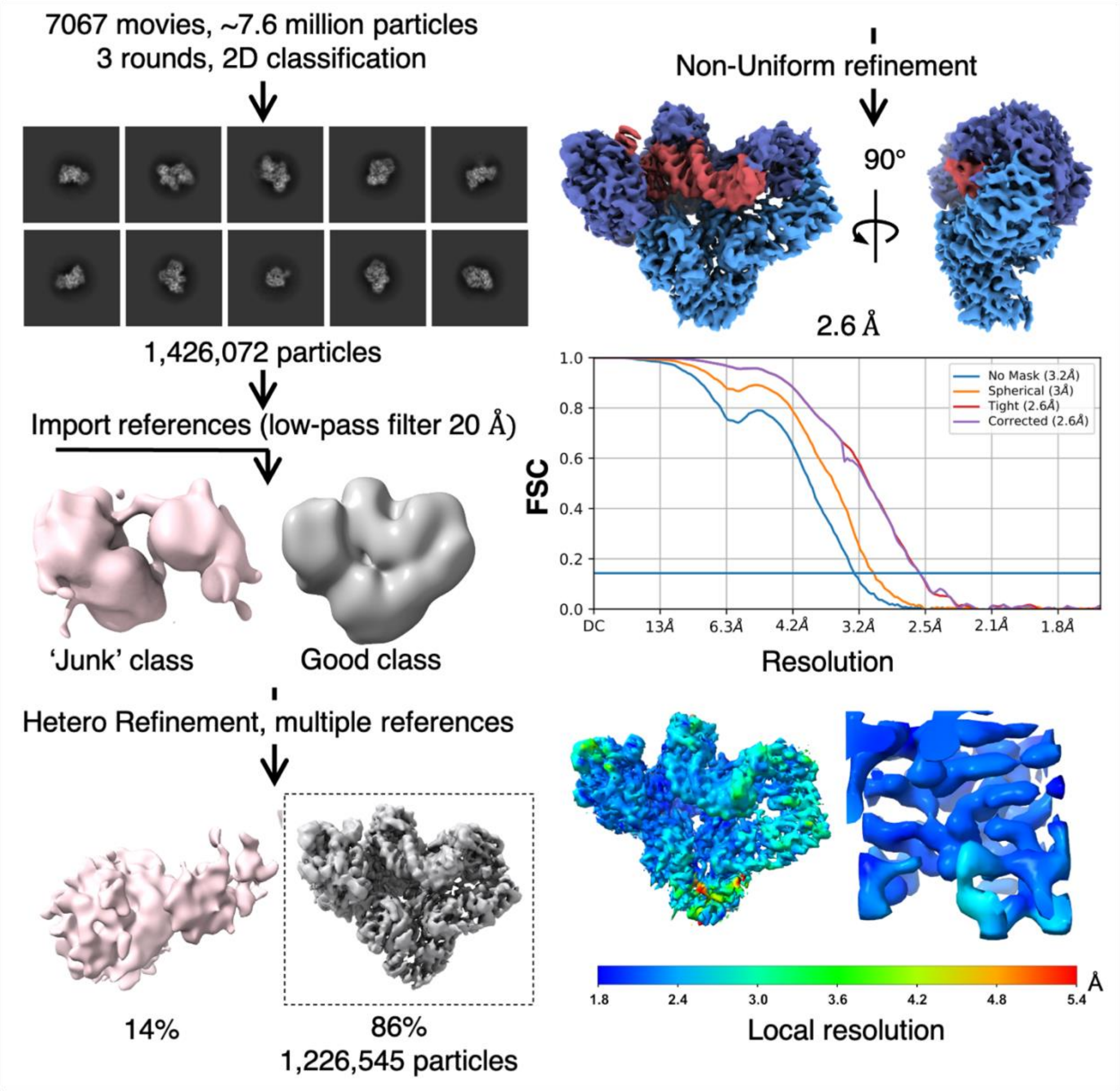


**Fig. S8.** Workflow of Cryo-EM processing of RT-DNA-dATP, State $P_{2min}$ (Data set 7). After 2D classification and 3D classification (Heterogeneous refinement), the clean particle stack is used for non-Uniform 3D refinement. All processing was done inside cryoSPARC pipeline. The resolution reported is based on the gold standard Fourier shell correlation (FSC) curve with cut-off value of 0.143.

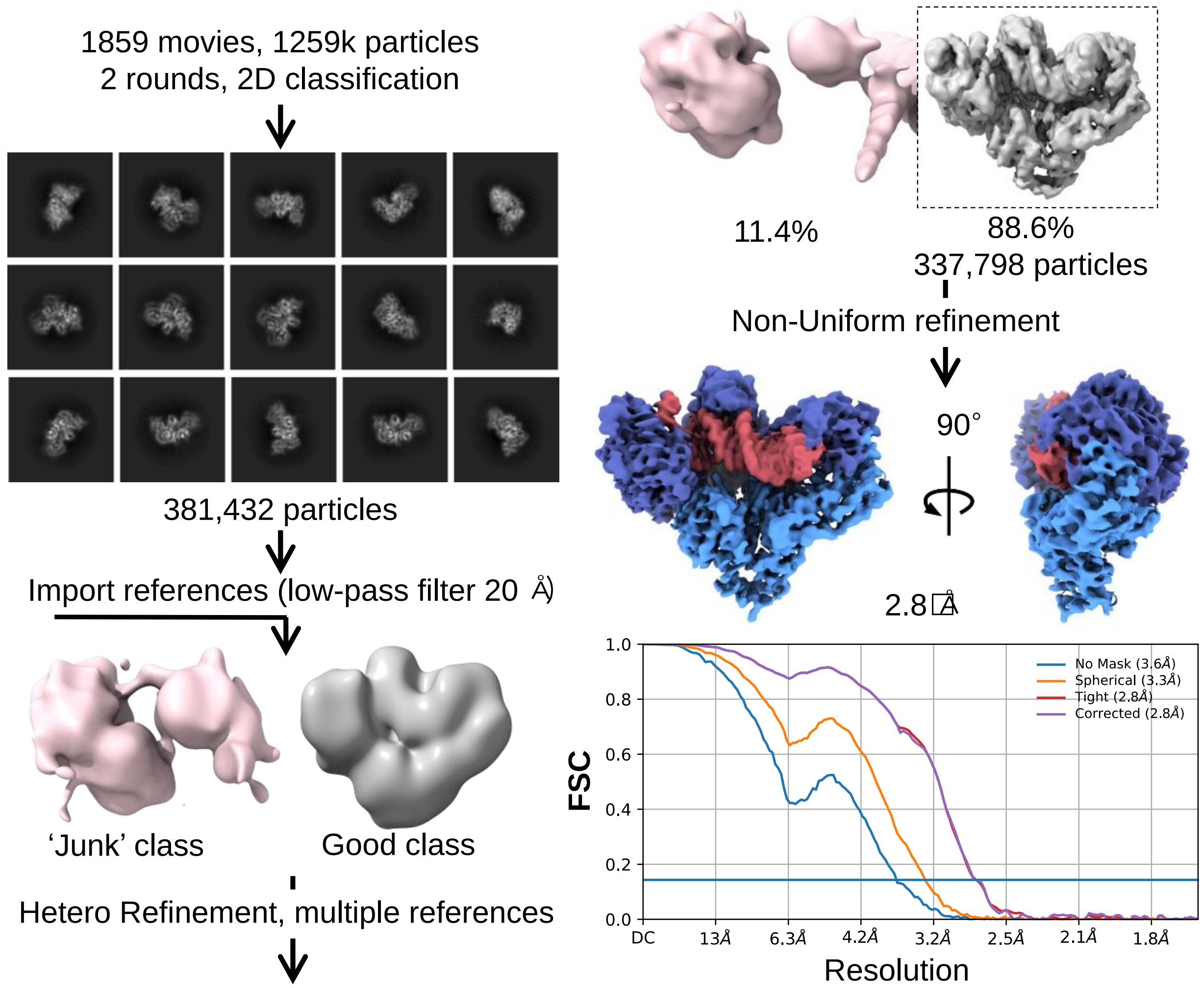


**Fig. S9.** Workflow of Cryo-EM processing of RT-DNA-dATP, State $P_{6min}$ (Data set 8). After 2D classification and 3D classification (heterogeneous refinement), the clean particle stack is used for non-Uniform 3D refinement. All processing was done inside cryoSPARC pipeline. The resolution reported is based on the gold standard Fourier shell correlation (FSC) curve with cut-off value of 0.143.

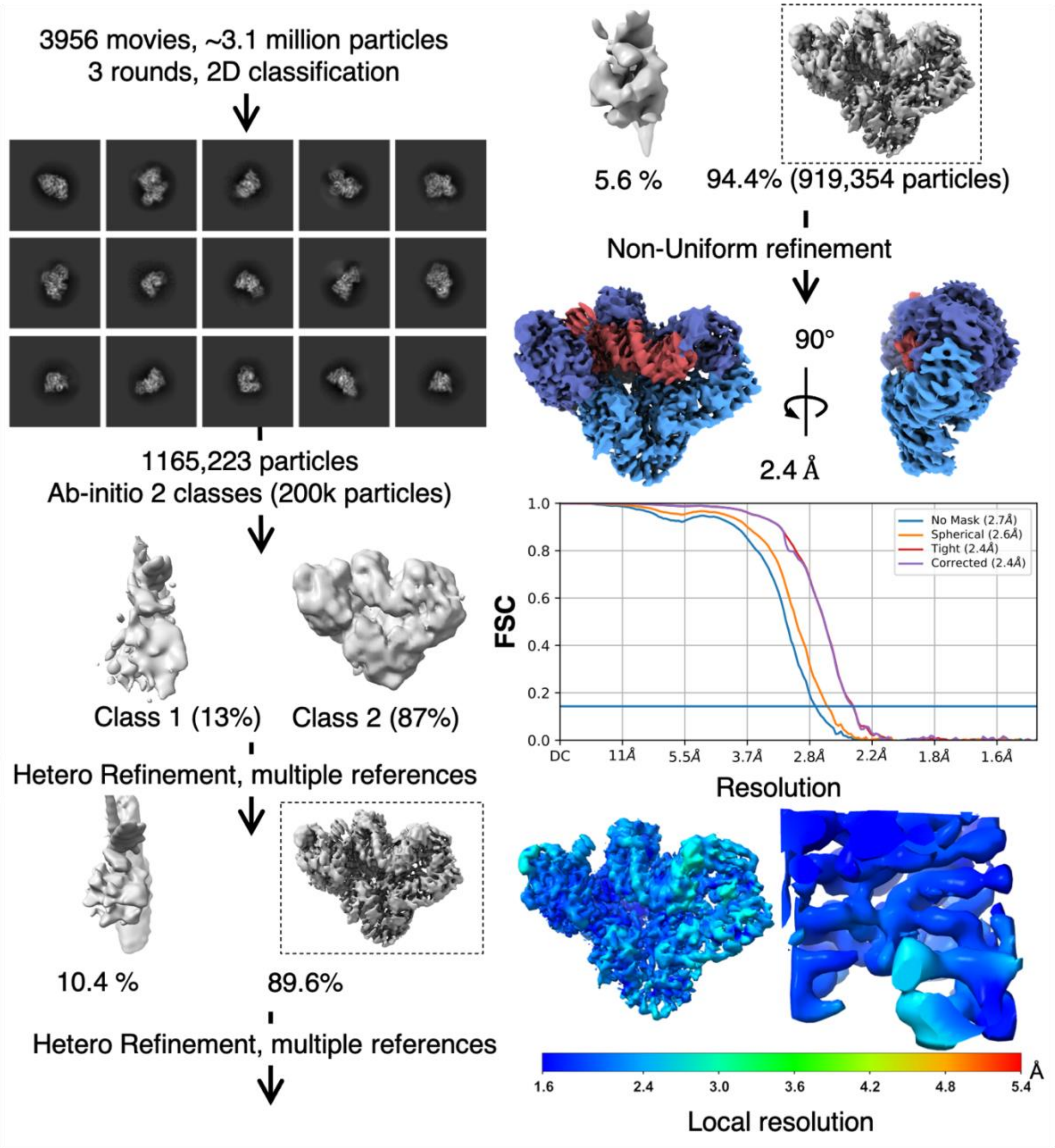


**Fig. S10.** Workflow of Cryo-EM processing of RT-DNA-dApCpp (Data set 9). After 2D classification and 3D classification, the clean particle stack is used for non-Uniform 3D refinement. All processing was done inside cryoSPARC pipeline. The resolution reported is based on the gold standard Fourier shell correlation (FSC) curve with cut-off value of 0.143.

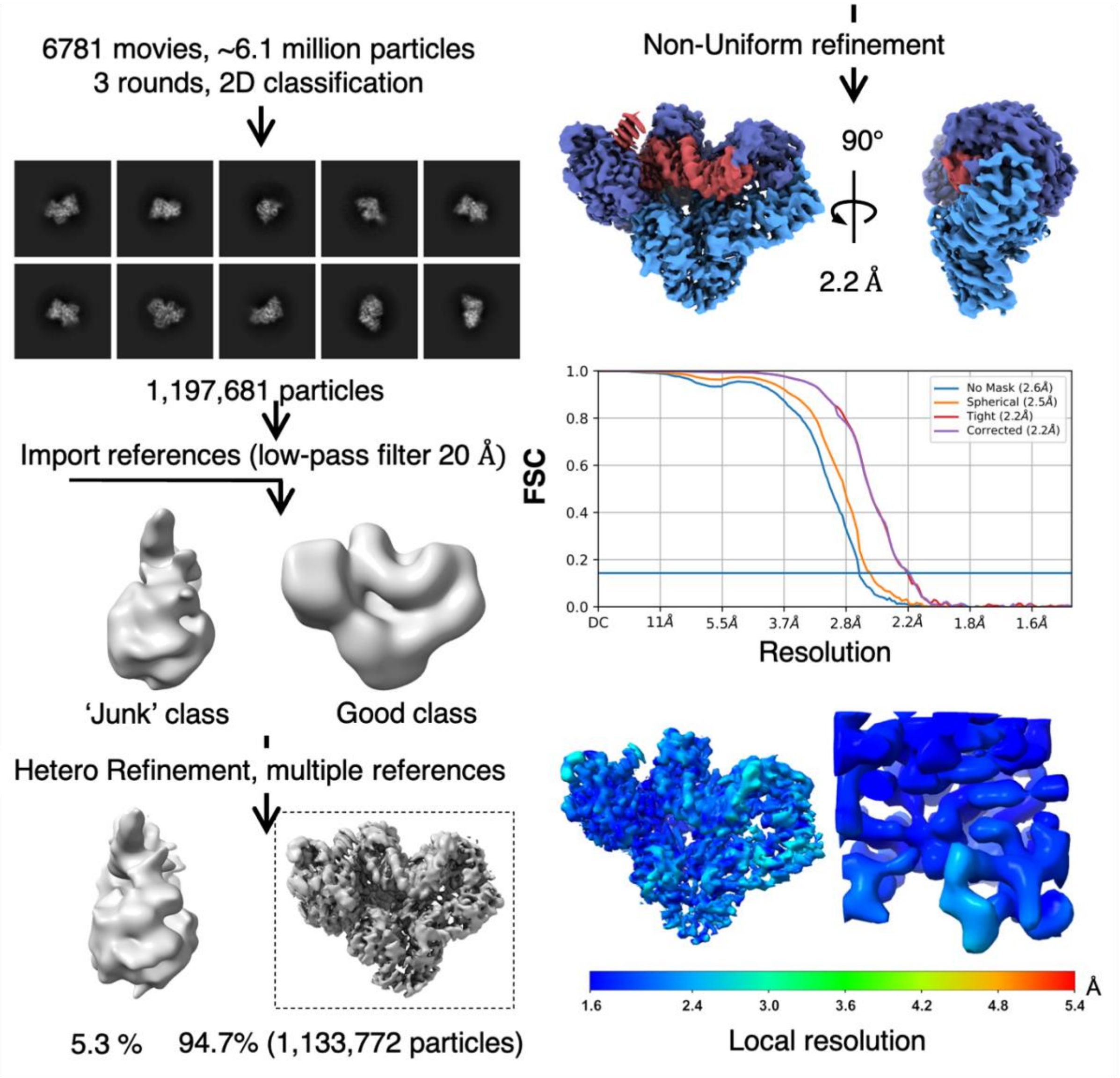


**Fig. S11.** Workflow of Cryo-EM processing of RT-DNA-dATP, $MgCl_2$ free (Data set 10). Reference classes for heterogeneous refinement are imported from data set 9 (Fig. S10). After 2D classification and 3D classification, the clean particle stack is used for non-Uniform 3D refinement. All processing was done inside cryoSPARC pipeline. The resolution reported is based on the gold standard Fourier shell correlation (FSC) curve with cut-off value of 0.143.

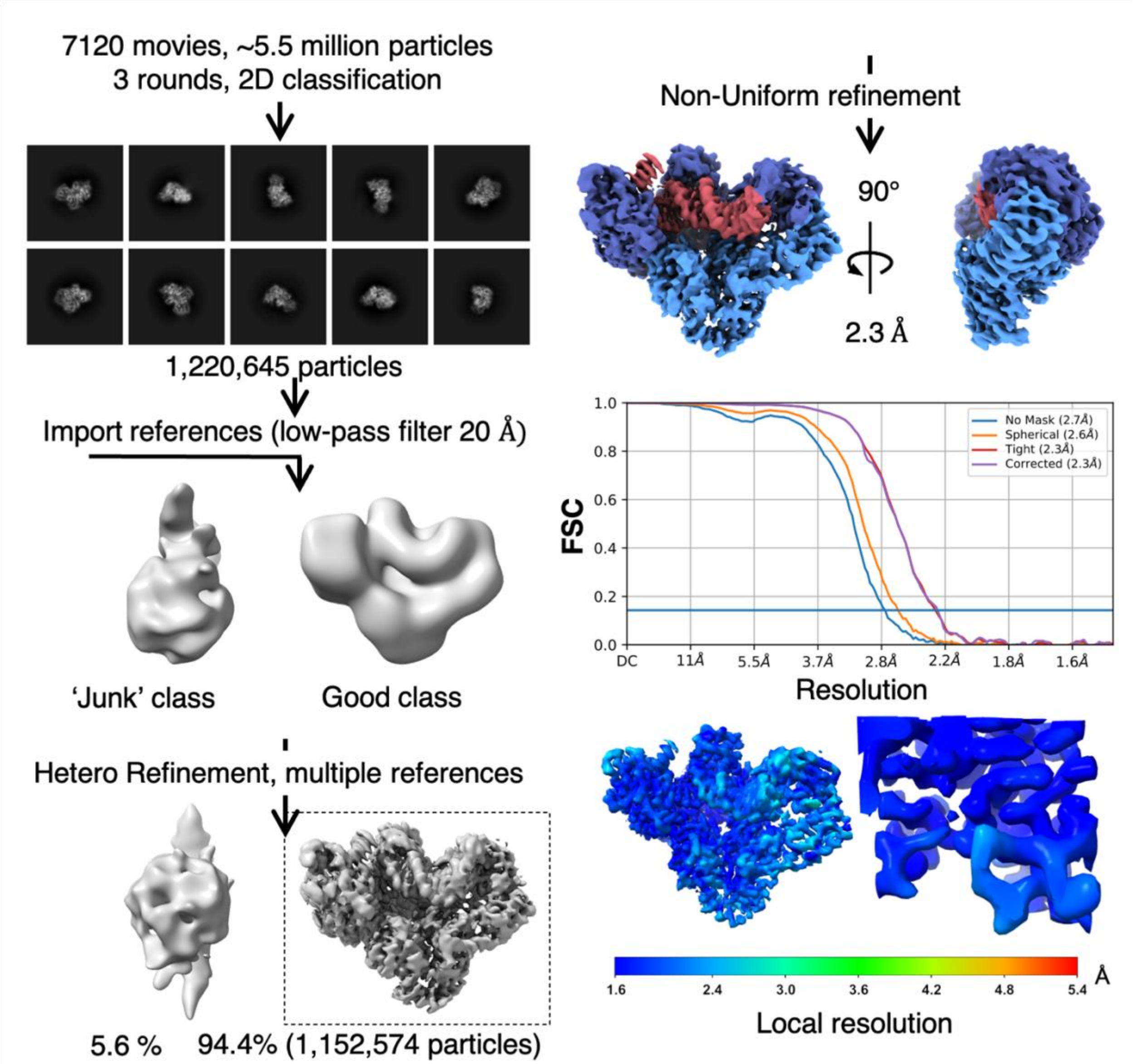


**Fig. S12.** Workflow of Cryo-EM processing of RT-DNA-dATP, 3'-terminated primer (Data set 11). Reference classes for heterogeneous refinement are imported from data set 9 (Fig. S10). After 2D classification and 3D classification, the clean particle stack is used for non-Uniform 3D refinement. All processing was done inside cryoSPARC pipeline. The resolution reported is based on the gold standard Fourier shell correlation (FSC) curve with cut-off value of 0.143.
